# Replication-coupled maturation of readthrough late transcripts by a polyomavirus small T antigen

**DOI:** 10.64898/2026.11.03.764058

**Authors:** Jeffery Nurdin, Amanda E. Moore, Kaitlyn Devine, Joseph Collura, Michelle Khattri, Masahiro Shuda

## Abstract

Productive polyomavirus gene expression requires the maturation of long, non-polyadenylated "wraparound" transcripts generated by late-strand readthrough of the polyadenylation signal on the circular genome, giving rise to structural protein mRNAs and microRNAs. In Merkel cell polyomavirus (MCV), small T (sT) antigen, along with large T (LT) antigen-mediated DNA replication, drives this process. Deletion of sT (MCVΔsT) attenuates late gene transcription, abolishes splicing, eliminates VP1 expression, and traps unspliced transcripts in the nucleus. Exogenous sT restored transcription and splicing in an aphidicolin-sensitive manner, requiring domains that activate DNA replication and bind chromatin. RNA sequencing revealed that sT promotes splicing and polyadenylation of wraparound transcripts, thereby increasing cytoplasmic VP1 and VP2 mRNA levels. A dual-fluorescence reporter confirmed that the leaky late poly(A) signal (LpA) was used more efficiently upon sT expression, indicating enhanced polyadenylation. Co-expression with LT abolished this effect, likely by overwhelming the capacity of the polyadenylation machinery. sT from HPyV7 and HPyV9 could similarly rescue this defect, indicating that this regulatory activity is conserved in a subset of sT proteins. Together, these findings define sT as a replication-coupled regulator of late transcript expression, splicing, and polyadenylation, revealing how a compact DNA virus converts readthrough transcription into productive gene expression.

## Introduction

Replication-coupled activation of late viral genes is a conserved regulatory strategy shared across diverse DNA virus families, including β- and γ-herpesviruses (1), adenoviruses (2), poxviruses (3), and baculoviruses (4), highlighting its broad importance in the DNA virus life cycle. However, the mechanistic link between genome replication and the transcriptional and post-transcriptional control of gene expression remains to be elucidated.

Polyomaviruses are small, circular double-stranded DNA viruses whose life cycle is governed by bidirectional early and late promoters within the non-coding control region (NCCR). The constitutively active early promoter drives the expression of the early gene, which is alternatively spliced to produce two conserved proteins: large T (LT) and small T (sT) antigen. LT is a replicative helicase that binds to the viral origin within the NCCR and unwinds the viral DNA through ATP-dependent activity. The role of sT varies among different polyomaviruses. Merkel Cell Polyomavirus (MCV) requires sT for continuous replication and viral gene activation, whereas BKV sT inhibits replication. The late promoter, induced only after viral DNA replication, drives expression of the late gene, which is similarly to early gene, spliced to produce the major and minor capsid proteins VP1 and VP2 (5,6).

In *betapolyomaviruses*, including SV40, JCV, and BKV, late gene activation is tightly associated with LT binding to the NCCR, where LT acts as a transcriptional activator (7-9). In murine polyomavirus (MPyV), an *alphapolyomavirus*, late gene activation is directly coupled to viral DNA replication (10). Replication disrupts late gene transcription termination on the circular genome, causing RNA polymerase to read through repeatedly and generate stable, multi-genomic “giant” transcripts that give rise to microRNAs (11,12) and are efficiently spliced into capsid mRNAs (10). While this readthrough phenomenon links structural late gene expression to regulatory miRNA production and appears to be conserved in polyomaviruses (13-15), the coordination of viral DNA replication, late gene activation, and 3′-end processing of late transcripts to drive life cycle progression remains unclear.

MCV, an *alphapolyomavirus* with a 5.4 kbp circular genome, causes most Merkel cell carcinoma (MCC) but is also a commensal skin virus that persists and sheds chronically from the skin of healthy individuals (16), where seroreactivity to its major capsid protein VP1 suggests ongoing cycles of viral DNA replication and virion production. However, the regulation of this life cycle remains poorly understood. Late genes are activated only after early gene transcription and DNA replication, and VP1 mRNA is generated exclusively through alternative splicing of the late transcript, which encodes the major capsid protein that self-assembles into virus-like particles. LT protein levels are further controlled by sT, which dampens Fbw7-mediated LT degradation via its LT stabilization domain (LTSD) (17). We previously showed that although sT is dispensable for initiating MCV DNA replication, an sT-deleted MCV mutant (MCVΔsT) cannot maintain its genome, fails to activate late genes, and does not produce VP1, indicating the pivotal role of sT in life cycle progression (5). An sT complementation study in MCVΔsT further demonstrated that late gene activation requires DNA replication. Unlike SV40, JCV, and BKV (7-9), and similar to MPyV (10), MCV late gene activation appears directly coupled to viral DNA replication rather than to LT-associated direct transcriptional activation.

In this study, we show that sT is a trans-acting regulator of late transcript maturation that directly promotes both splicing and 3′-end cleavage/polyadenylation at a leaky, readthrough-prone viral poly(A) signal, a function genetically separable from its established role in late gene transcription. Because this readthrough generates a wraparound pre-mRNA that matures into tandem leader exons through unconventional leader-to-leader splicing, MCV extends this paradigm to a compact, genetically tractable system for dissecting how transcriptional readthrough and 3′-end processing are coordinated in the cis.

## Materials and Methods

### MCV genome and plasmids

pSMART MCV_WT_ EcoRI has been previously reported (5). Briefly, the entire MCV sequence from the MCV-HF clone (GenBank accession no. JF813003) was ligated into the pSMART HC Amp vector (Lucigen) via its EcoRI site, which cuts the MCV genome once within the T-antigen. We also generated pSMART-LC-Amp, which contains a single BsrFI site in its multiple cloning site, and used it to clone the MCV-HF sequence via BsrFI, which cuts the MCV genome once in the late region of the genome. MCVΔsT was generated as described previously (5). Overlap extension PCR was used to generate pSMART MCVΔsT_TC_ EcoRI. Two DNA fragments containing double termination codons downstream of the LT splicing donor were amplified using the primer set AvrII-MCV-NCR_F33/sT. dTC_R31 and sT.dTC_F31/BamHI-R24, respectively. The introduced termination codons are highlighted in the primer sequences (S1 Table). After gel purification, equimolar amounts of the two fragments were mixed and fused by a second PCR using AvrII-MCV-NCR_F33 and BamHI-R24 primer pairs. The resulting fragment was cloned into pSMART MCV_WT_ EcoRI via the AvrII and BamHI sites, replacing the wild-type sequence with the mutant one. LL splicing acceptor (SA) mutants were produced by replacing the wild-type MCV fragment spanning the unique HpaI and AvrII restriction sites with a PCR fragment amplified using primers LL Acceptor mut_F58 and VP2-HpaI-F25. The consensus AG splice acceptor sequence was mutated to GG in the LL Acceptor mut primer (S1 Table). The pcDNA6 vectors expressing codon-optimized wild-type sT, LT, and sT mutants 91-95A, 90-94A, and 90-95A were previously described (5). Codon-optimized sT expression vectors were used for all sT complementation experiments in this study except for the FLAG-MCV sT construct in Fig. 6. We also generated the previously reported CxCxxC domain mutants in the codon-optimized MCV sT construct (18). CA, CF, and CAF sT mutants were generated using site-directed mutagenesis and overlap extension PCR. The CA and CF mutagenesis primers were STco_CAC_mut_F40/R40 and STco_CFC_mut_F40/R40, respectively (S1 Table). Overlap-extension PCR was performed using these internal primers together with the flanking primers MCV sT_F_DraI.BamHI and MCV sT_R_XhoI (S1 Table) at equimolar ratios of the DNA fragments. The CAF double mutant was generated by applying CA mutagenesis primers to the sequence-confirmed CF mutant, followed by overlap extension PCR using equimolar fragment ratios. To generate sT–mNeonGreen fusion constructs, we first deleted the initiation codon from pmNeonGreen-N1 (Addgene) using site-directed mutagenesis PCR (mNG-delIC-F27 and mNG-delIC-R27 primer set) to prevent potential alternative translation initiation downstream of the fusion. The resulting pmNeonGreen-dATG-N1 was digested with NheI and EcoRI to clone wild-type and mutant STs, which were amplified from pcDNA6 sT constructs using the sTcomut-NheI-S/sTcomut-EcoRI-AS primer pair. FLAG-tagged sT genes from the 12 non-MCV human polyomaviruses, previously cloned into a modified pcDNA6 vector (19), were PCR-amplified using pcDNA6 MCS.F23 and reverse primers designed against the sequence covering the termination codon of each sT gene (Table S1) and recloned into the same vector using KpnI and SacII sites to generate pcDNA6 FLAG-HPyV1-HPyV13 sT constructs (excluding MCV). To generate pcDNA6 FLAG-MCV sT, a FLAG-tagged wild-type MCV sT cDNA fragment was amplified from pSMART MCV_WT_ using FLAG-MCV_sT.F81 and MCV_sT.R51, digested with SalI and SacII, and cloned into pcDNA6 using XhoI and SacII. A polyadenylation reporter driven by the late MCV promoter was generated using overlap extension PCR. First, to produce the late MCV promoter-ZsGreen1 fragment, the full-length MCV NCCR and ZsGreen1 fragments were amplified using L-ZsG reporter_F/VP2.NCR-R60 and VP2.NCR-ZsG-F60/ZsG reporter_R, and fused using the L-ZsG reporter_F/ZsG reporter_R primer pair. The IRES2 and mScarlet fragments were amplified separately using the MCS-IRES2_F/IRES2_R and mScarlet_F/mScarlet_R primer sets, respectively. The three equimolar fragments were then mixed and fused by overlap-extension PCR using the L-ZsG reporter_F and mScarlet_R primer pair to generate the MCV late promoter-ZsG-IRES2-mScarlet cassette flanked by MfeI and AgeI sites at the 5’ and 3’ ends. After digestion, this fragment was cloned into MfeI/AgeI-digested pcDNA6/V5-His B (Invitrogen) to produce the pLMCV ZsG-MCS-mSca vector. The MCV sequence containing both late polyadenylation (LpA) signals was amplified using the primers NheI-LpA_F25 and XhoI-LpA_R25 from MCV-HF and cloned into pLMCV ZsG-MCS-mSca via NheI and XhoI sites to generate pLMCV ZsG-LpA-mSca. Mutated LpA fragments were generated using sequential overlap extension PCR. The first poly(A) signal (AATAAA) was mutated to AGGAGA by fusing two fragments amplified with pAsite L1m_F25/NheI-LpA and pAsite L1m_R25/XhoI-LpA from MCV-HF. Once confirmed, this mutated fragment served as the template for a second round of overlap-extension PCR to mutate the remaining AATAAA site into GATGAA. These mutations did not affect the antisense LT amino-acid sequence. The mutated fragments were cloned into pLMCV ZsG-MCS-mSca using the same restriction sites to produce pLMCV ZsG-LpAm-mSca. The BGH poly(A) sequence, used as a control, was amplified using the NheI-BGH pA_F/XhoI-BGH pA_R primer pair and cloned using the same strategy to generate pLMCV ZsG-BGH-mSca.

To generate a circular MCV genome, wild-type and mutant pSMART MCV-HF plasmids were digested overnight at 37°C with either EcoRI or BsrFI, followed by EcoRV for 5 h at 37°C. This reaction produces a full-length linear MCV-HF genome with self-ligatable EcoRI/BsrFI ends and a pSMART-LC-Amp vector fragment with EcoRV ends. The digested sample was purified using plasmid purification columns (Thomas Scientific) to obtain linearized MCV DNA fragments. Recircularization of the MCV-HF genome was performed using the T4 DNA Ligase system (NEB) overnight at 16°C at a low DNA concentration (2 ng/µL). The recircularized genome was purified using the same plasmid purification column, eluted with 10 mM Tris (pH 8.0), and used directly for transfection without removing the pSMART vector fragment from purified DNA. The MCVΔsT_CL_-VP1-mScarlet circular genome was generated by recombineering from pMC-MCV-VP1-P2A-mS, which expresses P2A-mScarlet fused to the C-terminus of the VP1 protein (20).

### Southern blot

Non-isotopic Southern blotting for MCV detection was established in our previous report (1). Briefly, cell pellets were lysed using lysis buffer (0.1 M NaCl, 10 mM Tris-HCl pH 8, 25 mM EDTA pH 8, 0.5% SDS) and treated with 100 μg/mL Proteinase K (Invitrogen) at 37°C overnight. After treatment with RNase A (Sigma), genomic DNA was extracted using phenol-chloroform, precipitated with NaOAc and ethanol, and the resulting pellet was dissolved in nuclease-free water (NFW). Ten micrograms of genomic DNA were digested with DpnI (NEB) and EcoRI-HF (NEB) overnight and fractionated on a 0.8% agarose/TBE gel. After capillary transfer onto a positively charged nylon membrane (Millipore), the digested DNA was UV-crosslinked to the membrane and hybridized with a biotin-labeled MCV probe overnight at 37°C. The hybridized membrane was sequentially washed with washing buffers and incubated with streptavidin-IR800 (LI-COR) for 1 h. The membrane was imaged using a ChemiDoc Imaging System (Bio-Rad).

### Cell lines, cell culture, transfection, and electroporation

Human embryonic kidney 293 (293) and neonatal human foreskin fibroblast human telomerase reverse transcriptase (BJ-hTERT) cells were maintained in DMEM (Corning) supplemented with 10 % fetal bovine serum (FBS) and 5 mL of non-essential amino acids (NEAA) (Gibco). The cells were maintained in a 37°C incubator with 5% CO_2_. Transfection was performed at a 1:3 ratio of total DNA in µg to Lipofectamine 2000 (Invitrogen) in µL, according to the manufacturer’s protocol, unless otherwise specified.

### Semi-quantitative RT-PCR and quantitative real-time RT-PCR were performed

Cell pellets were collected at designated time points, and RNA was extracted using a Qiagen RNA purification kit according to the manufacturer’s protocol. After RNA was quantified, a total of 1 µg of RNA was subjected to reverse transcription using the iScript^TM^ cDNA Synthesis Kit (Bio-Rad). The resulting cDNA was amplified using Q5 Hot Start DNA Polymerase (NEB) for semi-quantitative RT-PCR or PowerUp^TM^ SYBR Green Master Mix (Applied Biosystems) for qPCR. To detect late gene splicing, early gene splicing, and LL splicing by semiquantitative PCR, VP2.-118.F20/VP2.535.R20, panT.-180.25F/panT.766.21R, and VP2.-118.F20/d5415-a5308 BSP_R primer pairs were used (S1 Table). For the semi-quantitative 3′ rapid amplification of cDNA ends (RACE) experiment, 500 ng of total RNA, treated with TURBO DNase using the TURBO DNA-free™ Kit (Invitrogen), was reverse-transcribed with SuperScript IV (Invitrogen) using the 3′RACE_OligodT-linker primer. The resulting cDNA was subjected to semi-quantitative RT-PCR using the 3′RACE adaptor and KASI-VP1_F20 primer pair (S1 Table).

For the polyadenylation reporter assay with the ZsG-mScarlet bicistronic reporter construct, 1 µg of total RNA was purified into mRNA using the NEBNext Poly(A) mRNA Magnetic Isolation Module (NEB) following the manufacturer’s standard protocol. The final total elution volume (16 µL) was used for cDNA synthesis with the iScriptTM cDNA Synthesis Kit (Bio-Rad), and PowerUpTM SYBR Green Master Mix (Applied Biosystems) was used for quantitative RT-PCR using ZsG (ZsG.440_F21/ ZsG.569_R21) and mScarlet (mSca.1.F21/mSca.1.R21) primer pairs (S1 Table).

### Western blot

Cell pellets were lysed using RIPA buffer and sonicated. The resulting lysates were quantified using a DC Protein Assay Kit (Bio-Rad). Samples were mixed with sample buffer and 5% beta-mercaptoethanol and boiled at 95°C for 10 min. Proteins were transferred onto a nitrocellulose membrane using a Criterion Blotter or Trans-Blot Turbo System. The resulting membrane was blocked using 5% skim milk, probed with primary and secondary antibodies, and imaged using a ChemiDoc Imaging System (Bio-Rad). For subcellular protein fractionation, previously established protocols were applied with minor modifications (21,22). sT-transfected cells in a 10-cm dish were lysed in five volumes of CSK buffer (10 mM PIPES pH 6.8, 300 mM sucrose, 100 mM NaCl, 3 mM MgCl_2_, 1 mM EGTA) supplemented with 0.1% Triton X-100 and EDTA-free protease inhibitor (Roche), hereafter referred to as E1 buffer. After centrifugation, the soluble proteins were collected as the “cytoplasmic” fraction. The insoluble pellet was then lysed in 2 volumes of CSK buffer supplemented with 0.1% Triton X-100, 300 mM NaCl, and EDTA-free protease inhibitor. The resulting soluble fraction was designated the "nucleoplasm" fraction. To release DNA-bound proteins, the pellet was resuspended in five volumes of benzonase (Millipore)–E1 buffer and incubated at 37°C for 30 min. The remaining chromatin-bound proteins, including histones, were solubilized in 100 μL of a solution containing 2% SDS and 50 mM Tris-HCl (pH 7.5) with an EDTA-free protease inhibitor cocktail.

### Infectious virion production and infection

Virion production and infection were performed as previously described, with minor modifications (23). Briefly, 50% confluent 293 cells with inducible sT (293-TRE-sT) were transfected with 10 µg of recircularized MCV DNA in a 10 cm dish. During the 9-10 days period, cells were split when they reached full confluence, resulting in 3-4 confluent 15 cm dishes at the end of the propagation. Cell pellets were collected, lysed, and treated with 0.1% benzonase at 37°C overnight. After debris clarification, the supernatant was layered on top of OptiPrep^TM^ (Sigma Aldrich) with varying concentrations (29%, 33%, 39%, and 60%) and centrifuged at SW55Ti rotor 40,000 rpm for 5 hours. Viral titers were determined using quantitative real-time PCR (qPCR) with primer pairs targeting VP2 (VP2-535-R20 and VP2-336-F20), and pSMART MCV_WT_ EcoRI was used as a dilution standard.

For infection, 2 × 10^5^ BJ-hTERT cells were seeded in serum-free F12 and DMEM in a 1:1 ratio and supplemented with 1 mg/mL type IV Collagenase (Gibco), bFGF, EGF, and 3µM metalloproteinase inhibitor CHIR9901 (Selleckchem). A total of 1×10^5^ equivalents of viral genome/cell were added to each well. Two days post-infection, FCS was added to a final concentration of 20%, and the cells were incubated for four days.

### Lentivirus production and transduction

293FT cells were transfected with pLenti TRE sT (24), pCD/NL-BH*DDD, and pCAG-VSV-G plasmids. Three days after transfection, the supernatant was collected and clarified by centrifugation. Transduction was performed by adding the supernatant to the prepared cells with 6 µg/mL polybrene (Sigma). Two days after transduction, selection using 0.5 µg/mL puromycin (InvivoGen) was performed until the mock cells showed 100% cytotoxicity.

### Structure prediction

The HPyV sT structure was predicted using AlphaFold3 and visualized using PyMOL ver 3.1.8.

### Bioinformatic analyses

RNA sequencing was processed using RStudio version 2026.07.1+147 and R version 4.6.0 (2026-04-24), aligned using Rsubread, sorted, and indexed using Rsamtools, and bed graphs were generated using GenomicAlignments, Genomicranges, and rtracklayer packages. Gene tracks were visualized using the UCSD Interactive Genome Viewer with the MCV genome as a reference sequence (25). Genome transcript abundances were generated using bowtie2 2.5.5 for alignment, samtools ver 1.23.1 for sorting and indexing, and Salmon ver 2.1.1. The junction/half-leader ratio was calculated by aligning 40 bp of the junction sequence and 40 bp of the half-leader with a similar GC content using Python ver. 3.14.6.

## Results

### sT influences late-stage gene splicing and the production of VP1 capsid proteins

We previously demonstrated that the sT-deleted MCV mutant (MCVΔsT) exhibits decreased viral DNA replication and mRNA expression, fails to express VP1, and is fully rescued by exogenous sT, consistent with the essential role of sT in viral life cycle progression (5). Here, we generated two sT mutants: MCVΔsT_CL_, by cleaving the sT coding region, and MCVΔsT_TC,_ by introducing a termination codon (TC) downstream of the LT splicing donor (**Fig. 1A**, left panel). We examined late gene processing in wild-type MCV (MCV_WT_) and MCVΔsT_TC_ using semiquantitative RT-PCR with primers flanking the VP1 splicing junction (**Fig. 1A**, right panel). In MCV_WT_-transfected cells, VP1 mRNA (151 bp) remained low until day 6 post-transfection (p.t.), whereas VP2 expression, particularly the unspliced isoform (VP2u, 653 bp), was detectable by day 2 p.t. In contrast, no VP1 mRNA was detectable in MCVΔsT_TC_-transfected cells, and VP2u transcripts, although present, remained lower than those in MCV_WT_ (**Fig. 1B and Fig. 1C**). Increased VP1 splicing coincided with increased viral DNA replication in both MCVΔsT_TC_ and MCVΔsT_CL_-infected cells (**Fig. 1D, Fig. S1A, and Fig. S1B**) (5). In cells transfected with the replication-defective mutants MCV_rep(-)_ (26), VP2u transcripts were detectable by day 2 post-transfection but dissipated thereafter, and VP1 was not detected (**Fig. S1C**). Early gene splicing was unaffected by sT deletion, except for a modest decrease in unspliced T antigen transcript levels (**Fig. S1D and Fig. S1E**).

**Figure 1.**
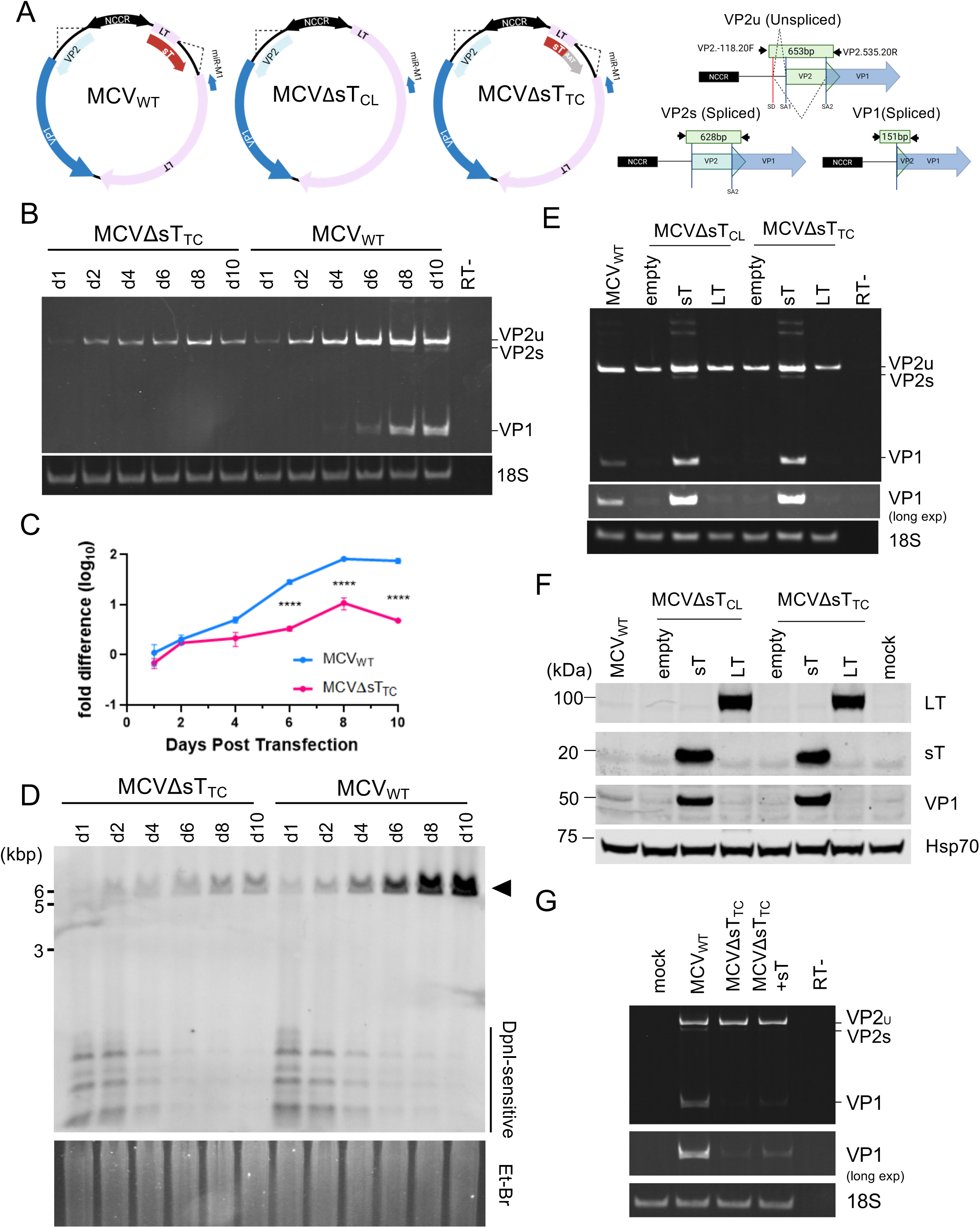
Late gene splicing is absent in sT-deleted mutants. **(A)** Schematic representation of the MCV constructs and late gene primer design used in this study. Unspliced late gene (VP2u), spliced VP2 (VP2s), and VP1 produced 653, 628, and 151 bp bands, respectively. **(B)** 293 cells were transfected with MCV_WT_ and MCVΔsT_TC_ and harvested at various time points for up to 10 days post-transfection. Total RNA was subjected to random hexamer RT-PCR to detect VP1 and VP2 RNA. No reverse transcription (RT-) was used as a negative control, and 18S rRNA was used as a loading control. **(C)** Samples from **B** were subjected to qRT-PCR, and statistical significance was determined using a two-way ANOVA. ****, p<0.0001 **(D)** DNA was extracted from the same sample of **B.** The cells were digested overnight with DpnI and EcoRI and subjected to Southern blotting using an MCV probe. The MCV genome is indicated by a closed arrowhead. **(E)** sT complementation experiments were performed by co-transfecting the MCV genome with empty, sT, and LT expression vectors. No RT was used as a negative control, and the 18S gene was used as a loading control. **(F)** Samples from **D** were lysed using lysis buffer, quantified, and subjected to western blotting. The membranes were probed with anti-LT, sT, VP1, and Hsc70 antibodies. **(G)** BJ-hTERT-TRE-sT cells were infected with MCV_WT_ or MCVΔsT_TC_ ± sT induction (MCVΔsT_TC_ only), harvested on day 7 (sT induced day 3), and total RNA analyzed by random-hexamer RT-PCR for VP1/VP2 expression.

To determine whether sT promotes late gene splicing through its role in stabilizing LT or independently of it, we co-transfected MCVΔsT_CL_ and MCVΔsT_TC_ with sT-or LT-expressing plasmids. Late gene splicing was restored when we overexpressed sT, but not LT, indicating that sT promotes splicing independent of its LT stabilization function (**Fig. 1E**). Western blotting confirmed that VP1 protein was detected only in the presence of exogenous sT, consistent with its requirement for efficient splicing and protein expression (**Fig. 1F**). Exogenously transfected sT and LT were readily detected, whereas endogenous levels were too low to be detected - LT alone in MCVΔsT-transfected cells, and both sT and LT in MCV_WT_-transfected cells.

MCV naturally infects dermal fibroblasts (27). To confirm this phenotype beyond 293 cells, we generated BJ-hTERT-TRE-sT cells with inducible sT and infected them with MCV_WT_ or MCVΔsT_TC_ viruses. Consistent with the 293-transfection model, MCV_WT_ exhibited late gene splicing, which was absent in MCVΔsT_TC_-infected cells. Inducing sT expression with doxycycline partially restored VP1 splicing in MCVΔsT_TC_ cells, but not to MCV_WT_ levels **(Fig. 1G)**. Together, these results indicate that sT is critical for late gene mRNA transcription and maturation, VP1 major capsid protein production, and, more broadly, the progression of the viral life cycle.

### Replication-associated domains of sT are important for late gene splicing

To identify the sT domain required for VP1 splicing, we examined previously reported mutants: an LTSD mutant (sT_90-94A_), an iron-sulfur-binding-defective mutant with two CxCxxC motifs mutated to AxAxxA (sT_CAF_), a DnaJ domain (sT_D44N_), and a PP2A-binding mutant (sT_L142A_) (18). qRT-PCR demonstrated that ST_90-94A_ and ST_CAF_ mutants failed to restore late gene splicing, whereas sT_D44N_ and sT_L142A_ restored it (**Fig. 2A**). Additional LTSD mutants (sT_91-95A_ and sT_90-95A_) and single CxCxxC mutants (sT_CA_ and sT_CF_) likewise failed to rescue VP1 splicing (**Fig. S2A**). Consistent with these RT-PCR results, VP1 protein expression in MCVΔsT_CL_ expression could only be rescued by DnaJ and PP2A-binding mutants, but not by LTSD and CxCxxC mutants (**Fig. S2B).**

**Figure 2.**
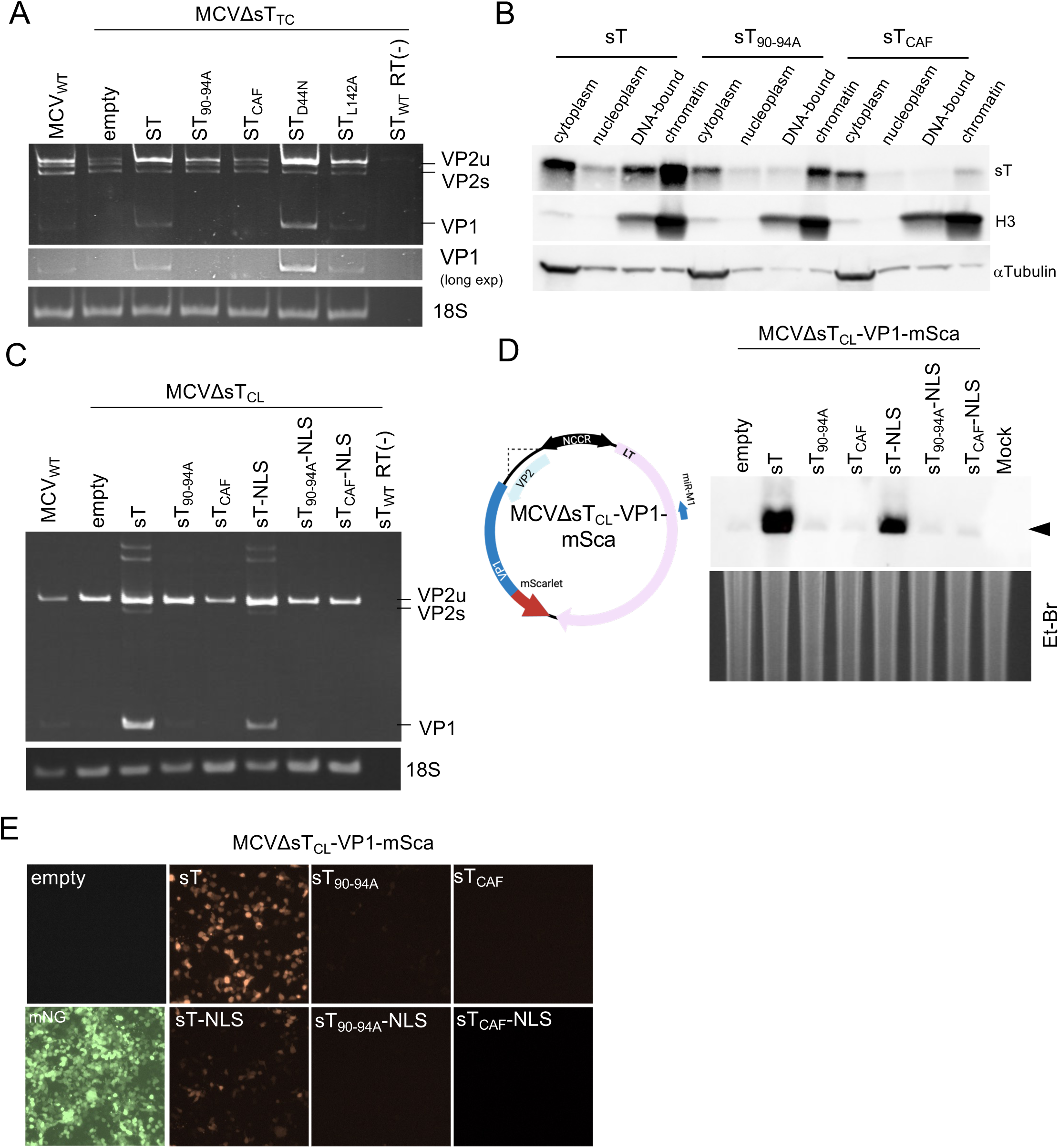
Viral DNA replication and sT replication-enhancing domains are important for late-gene splicing. **(A)** 293 cells were transfected with MCV_WT_ alone or with MCVΔsT_TC_ together with empty vector, sT_WT_, sT_90-94A_, sT_CAF_, sT_D44N_, or sT_L142A_. Total RNA was subjected to random hexamer RT-PCR. No RT (RT-) was used as a negative control, and 18S was used as a loading control. **(B)** 293 cells were transfected with sT, sT_LTSD_, or sT_CAF_, and the cell pellets were lysed and fractionated. The resulting fractions were subjected to western blotting. **(C)** 293 cells were transfected with MCV_WT_ alone or MCVΔsT_TC_ together with empty vector, sT, sT_90-94A_, sT_CAF_, sT-NLS, sT_90-94A_-NLS, and sT_CAF_-NLS. Total RNA was subjected to random hexamer RT-PCR. No RT was used as a negative control, and 18S was used as a loading control. **(D)** Schematic representation of MCVΔsT_CL_ with mScarlet-tagged VP1 (left panel). 293 cells were transfected with MCVΔsT_CL_-VP1-mSca and empty vector, sT, sT_90-94A_, sT_CAF_, sT-NLS, sT_90-94A_-NLS, or sT_CAF_-NLS. Genomic DNA was extracted, digested with EcoRI and DpnI, and subjected to Southern blotting. MCV genome was indicated by a closed arrowhead. **(E)** Visualization of mScarlet expression in samples shown in **D** using fluorescence microscopy. mNeonGreen (mNG) was used as a control for transfection efficiency.

Interestingly, sT_90-94A_-mNG and sT_CAF_-mNG were localized predominantly to the cytoplasm, whereas sT-mNG was localized to both the nucleus and cytosol, as previously reported (28) (**Fig. S2C**). Therefore, we tested whether LTSD and CxCxxC mutations altered the biochemical properties of sT using subcellular fractionation. Wild-type sT was detected predominantly in the cytoplasmic and chromatin fractions, with a small proportion in the DNA-bound fraction. Strikingly, sT_CAF_ lost its expression in the chromatin and DNA-bound fractions, and sT_90-94A_ also showed a significant decrease in DNA-bound fractions, whereas both mutants were abundantly expressed in the cytoplasmic fraction (**Fig. 2B**). Because sT lacks a canonical nuclear localization signal (NLS), we predicted that its nuclear localization is mediated by its DNA and chromatin-binding activity, and that the LTSD and CxCxxC mutations exclude sT from the nucleus indirectly by disrupting chromatin tethering. Consistent with this, fusing the SV40 LT NLS into MCV sT restored the nuclear expression of sT_90-94A_ and sT_CAF_ proteins (**Fig. S2C**). However, sT_90-94A_-NLS and sT_CAF_-NLS still failed to rescue VP1 splicing in MCVΔsT_CL_, although sT-NLS activity in inducing VP1 splicing was less efficient than that of wild-type sT. These data indicate that forcing nuclear entry through a heterologous NLS cannot substitute for chromatin-binding activity and that VP1 splicing requires the chromatin/DNA-binding function of the LTSD and CxCxxC motifs, rather than nuclear presence alone (**Fig. 2C and S2D**).

sT promotes LT-mediated viral origin replication through LTSD and CxCxxC motifs (5,18,28). We tested the effects of these mutants on DNA replication by complementing MCVΔsT_CL_-VP1-mSca, an MCVΔsT_CL_ reporter genome carrying a C-terminally mScarlet-tagged VP1, with each sT construct. Consistent with previous results, Southern blotting showed that only sT and sT-NLS supported DNA replication, whereas sT_90-94A_ and sT_CAF_ did not, regardless of NLS tagging (**Fig. 2D and Fig. S2E**). Accordingly, mScarlet expression was observed only in cells complemented with sT- and sT-NLS, with no signal in cells complemented with other sT mutants (**Fig. 2E**). Inhibiting DNA replication by aphidicolin similarly ablated VP1 and VP2 splicing, leaving VP2 in its unspliced form (**Fig. S2F**). Taken together, these results indicate that late gene splicing requires both replication-associated domains of sT and active DNA replication.

### sT promotes late transcripts maturation

Because sT deletion ablates VP1 splicing and protein expression, we investigated whether it also affects the nuclear-cytoplasmic distribution of late transcripts. We transfected sT-inducible 293 cells (293-TRE-sT) with MCV_WT_ or MCVΔsT_TC_, with or without sT complementation, and fractionated the nuclear and cytoplasmic RNA. In MCV_WT_ and MCVΔsT_TC_+sT, the nuclear fraction showed VP2u and VP1, whereas the cytoplasmic fraction showed reduced VP2u levels but was enriched with spliced VP1. In contrast, MCVΔsT_TC_ alone showed nuclear VP2u but no spliced VP1 in either fraction, indicating a failure to produce mature VP1 mRNA (**Fig. S3A**). Infection of BJ-hTERT-TRE-sT fibroblast cells with MCV_WT_ or MCVΔsT_TC_ recapitulated this pattern: spliced VP1 was detected in the cytoplasmic fraction of MCV_WT_ and MCVΔsT_TC_+sT, with nuclear VP1 undetectable in both, whereas MCVΔsT_TC_ alone showed no detectable VP2u or VP1 in either fraction **(Fig. S3B)**. qRT-PCR showed that late transcripts were predominantly nuclear across all constructs, but nuclear retention was higher for MCVΔsT_TC_ alone than for MCV_WT_ and MCVΔsT_TC_ + sT (**Fig. S3C and Fig S3D**). As expected, β-actin control transcripts were predominantly localized in the cytoplasm.

To further investigate the role of sT in late transcript maturation and localization, we performed short-read RNA sequencing in MCV-transfected 293-TRE-sT cells using total RNA with ribosomal RNA-depleted or poly(A)-enriched mRNA libraries (**Fig. 3A**). Cytoplasmic mRNA from MCV_WT_ and sT-rescued MCVΔsT_CL_ consisted predominantly of late gene transcripts with a small proportion of early genes, whereas MCVΔsT_CL_ mRNA without sT complementation demonstrated the opposite pattern, predominantly early transcripts with minimal late transcripts (**Fig. 3B**), indicating that sT is required for the early to-late switch in the MCV life cycle. To assess maturation, we compared the cytoplasmic mRNA (output) to the total nuclear RNA (input) within each sample. The early gene representation was similar between mRNA and total RNA across all samples. For late genes, however, the mRNA-to-total RNA proportion was highest in MCV_WT_, followed by MCVΔsT_CL_ + sT, and then MCVΔsT_CL_ (**Fig. 3C**). The resulting maturation ratio (mRNA/total RNA read coverage) demonstrated that early transcripts (57kT and LT) were largely unaffected by sT deletion, remaining at 200-400% across constructs. However, late gene maturation dropped sharply with sT loss: 40-48% of MCV_WT_ late transcripts were mature, versus only 3-4.5% for MCVΔsT_CL_ (**Fig. 3D**). sT complementation restored maturation in MCVΔsT_CL_+ sT to only 8.5-12%, still below MCV_WT_ levels, likely because sT-enhanced transcription outpaced splicing/maturation capacity (Fig. 4C). A similar result was observed for MCVΔsT_TC_ (**Fig. S4A-C**). Together, these findings underscore sT’s importance of sT in late gene RNA maturation during the MCV life cycle.

**Figure 3.**
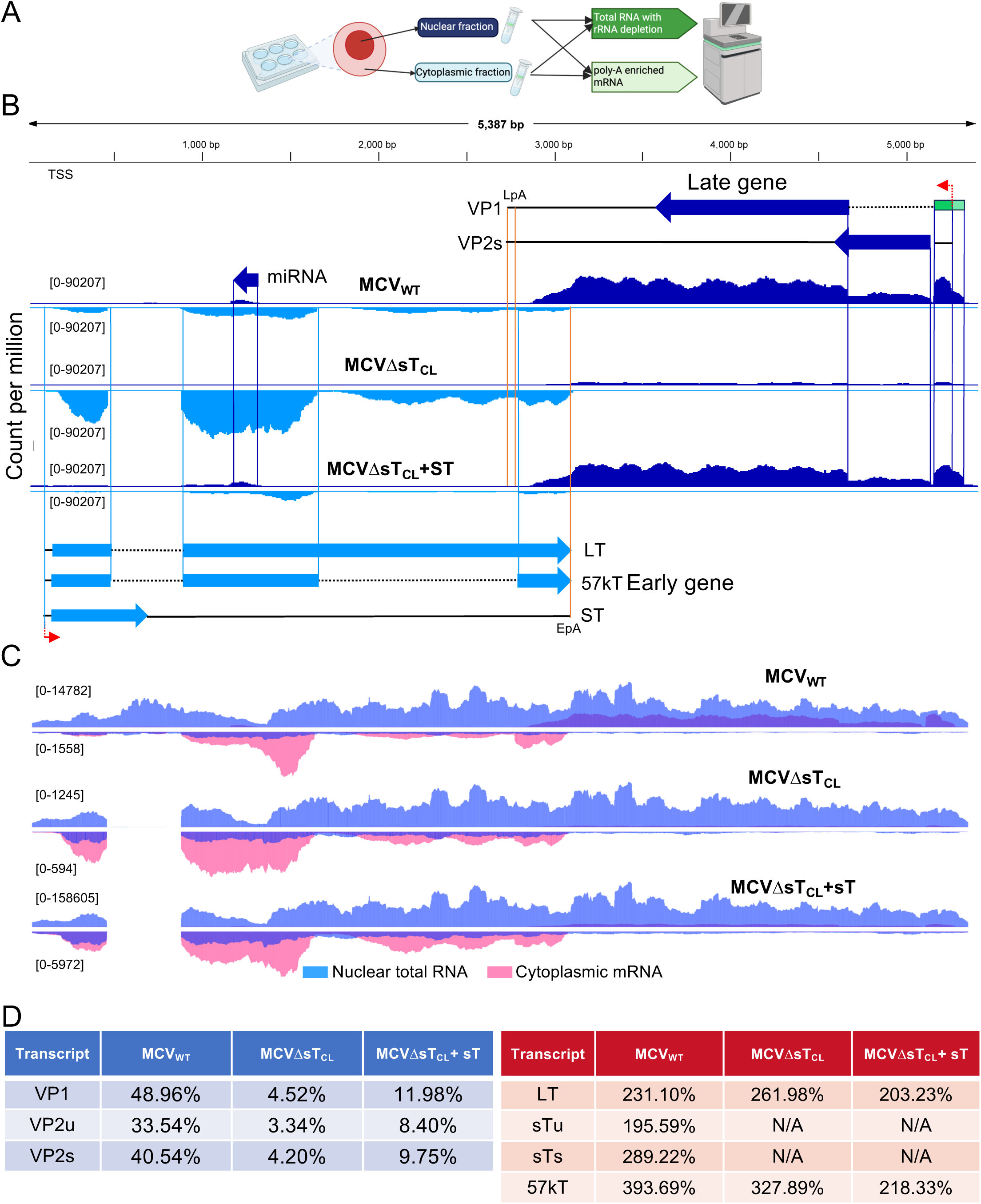
sT drives late gene RNA export and early-to-late switch. **(A)** Schematic representation of RNA sequencing preparation. Nuclear and cytoplasmic fractions were separated, and each fraction was subjected to RNA sequencing using either ribosomal RNA depletion (total RNA-seq) or poly(A) enrichment (mRNA-seq). (**B)** Normalized read coverage of poly(A)-enriched RNA-seq from the cytoplasmic fraction of MCV_WT_, MCVΔsT_CL_, and sT(codon-optimized)-complemented MCVΔsT_CL_. Late genes are represented at the top in dark blue, and early genes are represented at the bottom in light blue. Boxes and arrows represent coding sequences; bold lines represent the exonic regions, and dashed lines represent the intronic regions. Green boxes represent a leader exon. Red lines with arrows represent transcription start sites. EpA, early poly(A); LpA, late poly(A). Only representative proteins are shown because of space constraints. Numbers signify y-axis range read counts. **(C)** Read coverage of poly(A)-enriched RNA-seq of the cytoplasmic fraction (pink) and ribo-depleted total RNA-seq of the nuclear fraction (purple) of MCV_WT_, MCVΔsT_CL_, and sT-complemented MCVΔsT_CL_. Late genes are represented at the top and early genes at the bottom of the graph. Each graph was scaled independently. Numbers signify y-axis range read counts. **(D)** Maturation rates from MCV_WT_, MCVΔsT_CL_, and sT-complemented MCVΔsT_CL_ were determined by dividing the of poly(A)-enriched RNA-seq count with rRNA-depleted total RNA count, and are presented as percentages.

**Figure 4.**
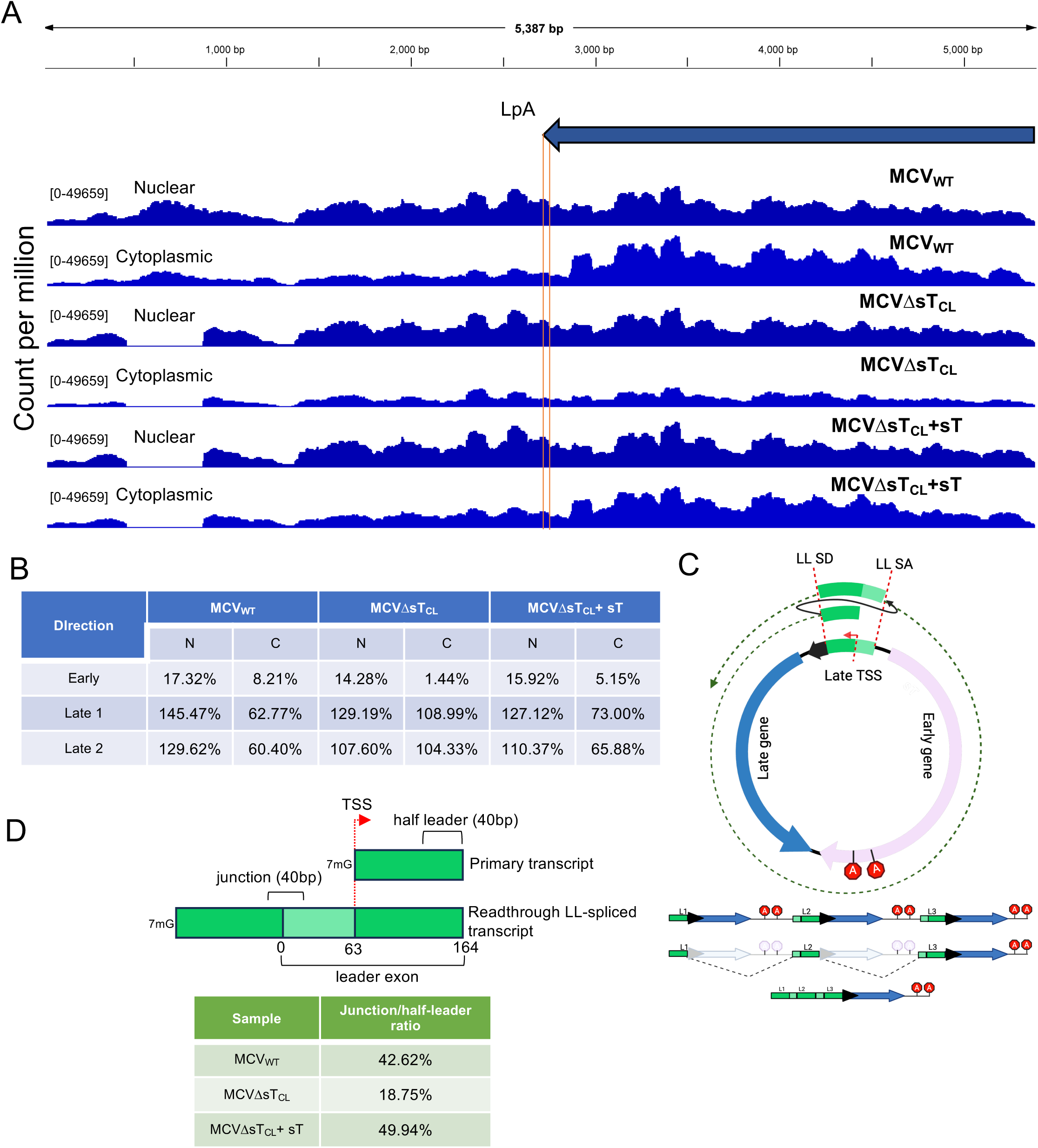
sT hinders LpA readthrough and promotes LL splicing. **(A)** Normalized read coverage of late genes from ribo-depleted total RNA-seq in MCV_WT_, MCVΔsT_CL_, and MCVΔsT_CL_ + sT. The LpA is indicated by a red line. Numbers signify the y-axis range read counts. **(B)** Readthrough ratio (RR) calculated by dividing post-poly(A) count with pre-poly(A) count, presented in percentage. N=nucleus, C=cytoplasm. **(C)** Schematic representation of LL-splicing in MCV. Late genes read through the polyadenylation signals and wrap around the transcription start site (TSS). The nascent transcript is spliced from each downstream SD back to the upstream SA, fusing repeated leader exons with each wrap-around cycle; this continues until the transcript reaches the late gene coding sequence, which is then polyadenylated, yielding a compact mRNA with multiple leader exons. **(D)** Schematic representation of the LL-splicing junction and half-leader sequence calculation method. Twenty nucleotides flanking the junction (pre- and post-junction) were used, and forty nucleotides with similar GC content from the half-leader sequence were selected as the denominator (top panel). Junction-to-half-leader read ratios from cytoplasmic mRNA-seq are presented as percentages.

### sT suppresses late gene readthrough

Total RNA sequencing has demonstrated that late gene transcripts read through late polyadenylation (LpA) sites in the nuclei of MCV_WT_-, MCVΔsT_TC_-, and MCVΔsT_CL_-transfected cells, although the readthrough is minimal for early genes (**Fig. S5A**). In MCV_WT_, MCVΔsT_TC_+sT, and MCVΔsT_CL_+sT, these post-LpA reads decreased markedly in the cytoplasm, indicating that readthrough transcripts are cleaved and polyadenylated at the LpA before nuclear export (**Fig. 4A and Fig. S5B**). In contrast, cytoplasmic post-LpA reads in MCVΔsT_TC_ and MCVΔsT_CL_ remained comparable to pre-LpA levels, indicating that in the absence of sT, late gene transcripts failed to terminate at the LpA.

To quantify this, we calculated the readthrough ratio (RR: post-LpA/pre-LpA read coverage) from total RNA-seq data. Early genes showed a low cytoplasmic RR (1-8%), consistent with the gene track in **Fig. S5A**. For MCV_WT_ late genes, the RR was high and variable in the nucleus (129-145%) but dropped to 60-62% in the cytoplasm, confirming that readthrough transcripts are processed before export. This processing was largely absent in MCVΔsT_CL_, where the RR remained high in both the nucleus (107-129%) and cytoplasm (104-109%). In MCVΔsT_CL_+sT, sT complementation restored readthrough termination, reducing the RR from 110-127% in the nucleus to 66-73% in the cytoplasm (**Fig. 4B**). A similar trend was observed for the MCVΔsT_TC._ (**Fig. S5C**).

Polyomaviruses process this giant wraparound transcript through a distinct mechanism called leader-to-leader (LL) splicing(14,15,25,29). MCV’s leader exon spans the late transcription start site, so the wraparound transcript undergoes LL splicing, joining the leader donor (LD) at the 3′ end of the half-leader exon to the leader acceptor (LA) at the 5′ end of the full-leader exon and excising the intervening sequence (**Fig. 4C**). Because the full-leader exon, and thus LA, forms only when RNA polymerase II reads through the genome without reinitiating, LL splicing and tandem leader exons occur exclusively in readthrough transcripts. Using mRNA sequencing, we assessed sT’s effect on LL splicing of these transcripts (**Fig. 4D**, top panel). In the presence of sT, 43-50% of half-leader exon reads (common to all late transcripts, regardless of readthrough status) from MCV_WT_ and MCVΔsT_CL_ + sT contained an LL splice junction, indicating efficient resolution of these giant wraparound transcripts into leader-to-leader spliced products. In the absence of sT, MCVΔsT_CL_ showed only 18% LL junction reads (**Fig. 4D**, bottom panel), indicating that splicing efficiency, not substrate abundance, limits LL splicing without sT. Together, these results indicate that sT is important not only for late gene body splicing but also for LL splicing of readthrough transcripts.

### sT promotes polyadenylation of late transcript

To measure the termination efficiency of MCV LpA, we generated a dual reporter (30) in which ZsGreen (ZsG), driven by the CMV promoter, is followed by LpA, an EMCV IRES, mScarlet, and a BGH poly(A) signal, such that mScarlet is expressed only upon readthrough past LpA (**Fig. S6A**). As controls, we replaced LpA with a mutant lacking the canonical AATAAA sequence (LpA(-)), which abolishes termination, or with a strong canonical BGH poly(A) signal (**Fig. S6A**). The wild-type LpA reporter showed a weak mScarlet signal, indicating that LpA is leaky and supports translation beyond LpA. The LpA(−) showed a stronger mScarlet signal, whereas the BGH poly(A) construct showed no signal, reflecting efficient termination (**Fig. S6A**). Quantification of the normalized mScarlet to ZsG fluorescence confirmed that the LpA(−) reporter produced the strongest readthrough signal (**Fig. S6A**).

To determine the role of sT in the polyadenylation of late transcripts, we replaced the CMV promoter with the MCV late promoter in the dual reporter system. We performed qRT-PCR using poly(A)-enriched mRNA from the reporter-transfected cells to quantify terminated monocistronic ZsG transcripts and readthrough dicistronic ZsG-mScarlet transcripts, calculating the readthrough efficiency as the mScarlet/ZsG ratio. sT expression reduced mature mScarlet mRNA levels, suggesting that sT promotes termination/polyadenylation at the leaky LpA site, whereas LT alone had no effect on mScarlet expression. Unexpectedly, however, this sT-mediated reduction was lost when sT was co-expressed with LT (**Fig. 5A**). As controls, the LpA(-) reporter again showed 3-fold higher read-though efficiency than wild-type LpA, whereas the BGH poly(A) reporter showed only about 25% of wild-type LpA’s read-through efficiency (**Fig. 5A**). Together, these results suggest that sT possesses a distinct polyadenylation-promoting role, independent of its role in enhancing promoter activity.

**Figure 5.**
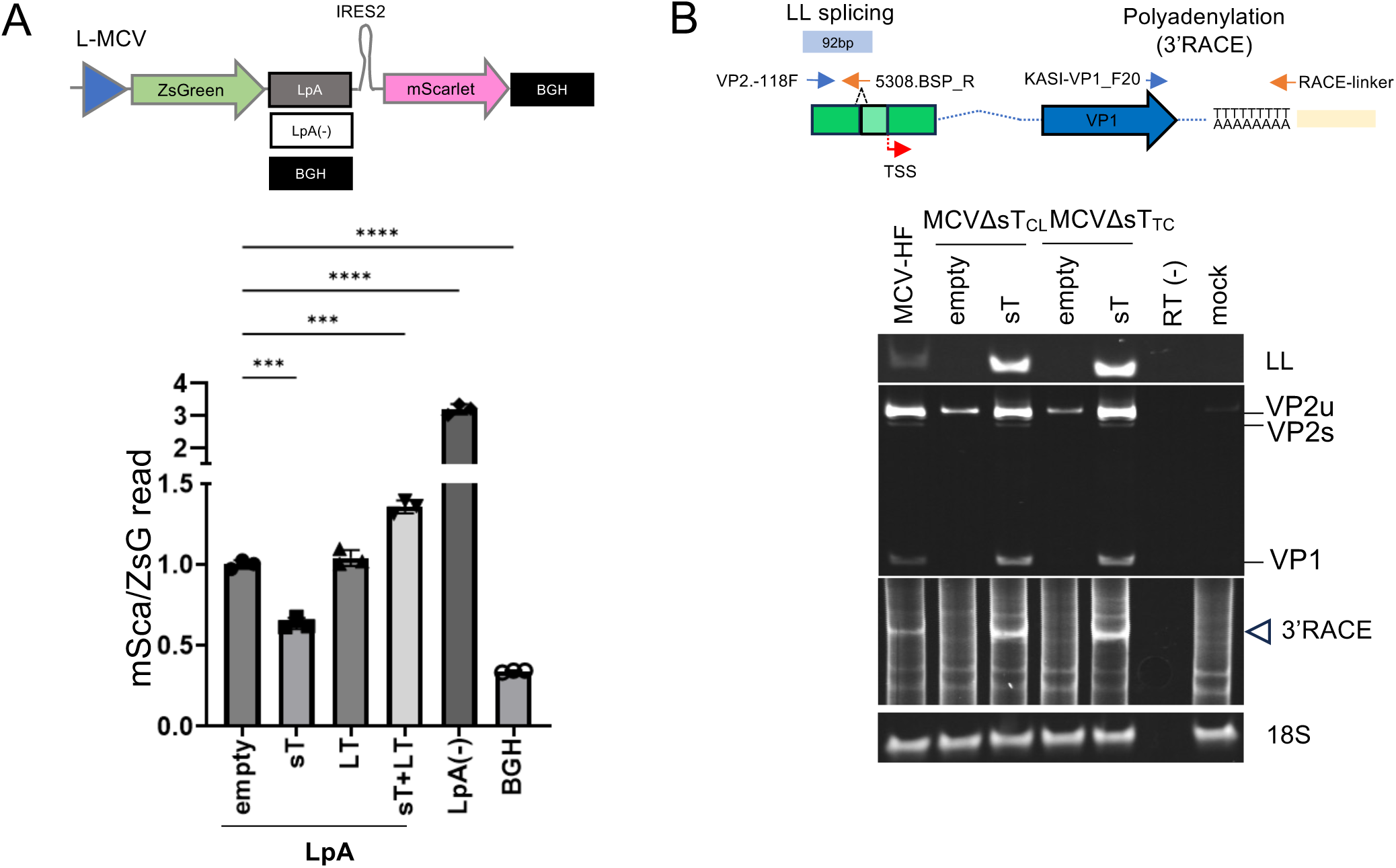
sT promotes polyadenylation of LpA. **(A)** Schematic representation of the dual-reporter system used in this assay (top panel). 293 cells were transfected with a dual reporter construct and an empty vector, sT, LT, or sT/LT, and total RNA was enriched for poly(A) before being subjected to qRT-PCR using ZsG or mScarlet-specific primer pair. mScarlet transcript abundance was normalized to ZsG transcript abundance using the 2^-ΔΔCt^ method. **(B)** Strategies and primers used to detect LL splicing and poly(A) tails. 293 cells were transfected with MCV_WT_, MCVΔsT_CL_, or MCVΔsT_TC_ together with an empty or sT-expressing vector. Total RNA was subjected to random hexamer RT-PCR to detect LL splicing and VP1/VP2 RNA, or to 3’RACE (oligo-dT-linker reverse transcription followed by PCR with a late-gene-specific primer) to detect the 3’ ends of polyadenylated late transcripts. No reverse transcription (RT-) was used as a negative control, and 18S rRNA was used as a loading control.

To determine whether sT similarly promotes polyadenylation and LL splicing in the native viral genome, we performed 3-RACE and LL junction-RT-PCR (25). Both were detected in MCV_WT_-transfected cells but not in MCVΔsT_CL_ or MCVΔsT_TC_, and both were restored by exogenous sT, indicating that sT promotes polyadenylation and LL splicing of the viral late transcripts, consistent with the higher LL splicing rate determined by RNA sequencing (**Figs. 5B and 4D**). Mutating the LL splice acceptor abolished LL splicing but did not prevent late gene splicing, regardless of sT complementation in MCV_WT_-transfected 293 cells. These results indicate that LL and VP1 splicing occur independently, although both are activated by sT (**Fig. S7C**).

### HPyV7 sT and HPyV9 sT showed similar phenotype to MCV sT

To investigate whether this late-gene splicing function is shared by other HPyVs, we cloned sT genes from 13 human polyomaviruses (HPyVs), including wild-type MCV sT as a control, and co-transfected each with MCVΔsT_TC_. We found that HPyV7 and HPyV9 sT were the only proteins that rescued late gene splicing in MCVΔST_TC_ despite AlphaFold3-predicted structural similarity among all 13 STs (**Fig. 6A and Fig. S7A)**. Quantification of late gene expression by qRT-PCR demonstrated that MCV, HPyV7, and HPyV9 sT all significantly increased transcription relative to uncomplemented MCVΔsT_TC_ without any complementation (**Fig. 6B**). Unlike HPyV7 and HPyV9, the HPyV6 sT restored transcription to MCV_WT_ levels without rescuing VP1 splicing, dissociating its transcriptional and splicing functions. While the LTSD domain is not conserved across sT proteins, the CxCxxC motif is conserved (**Fig. S7B**). Therefore, we generated CxCxxC mutants of HPyV7 and HPyV9 sT to determine whether this motif affects late gene splicing. As with MCV sT, CxCxxC mutations in HPyV7 and HPyV9 sT cause loss of VP1 splicing and decreased late gene transcription, suggesting that this motif is critical for both late gene transcription and splicing in these HPyVs (**Fig. 6C and Fig. 6D**).

**Figure 6.**
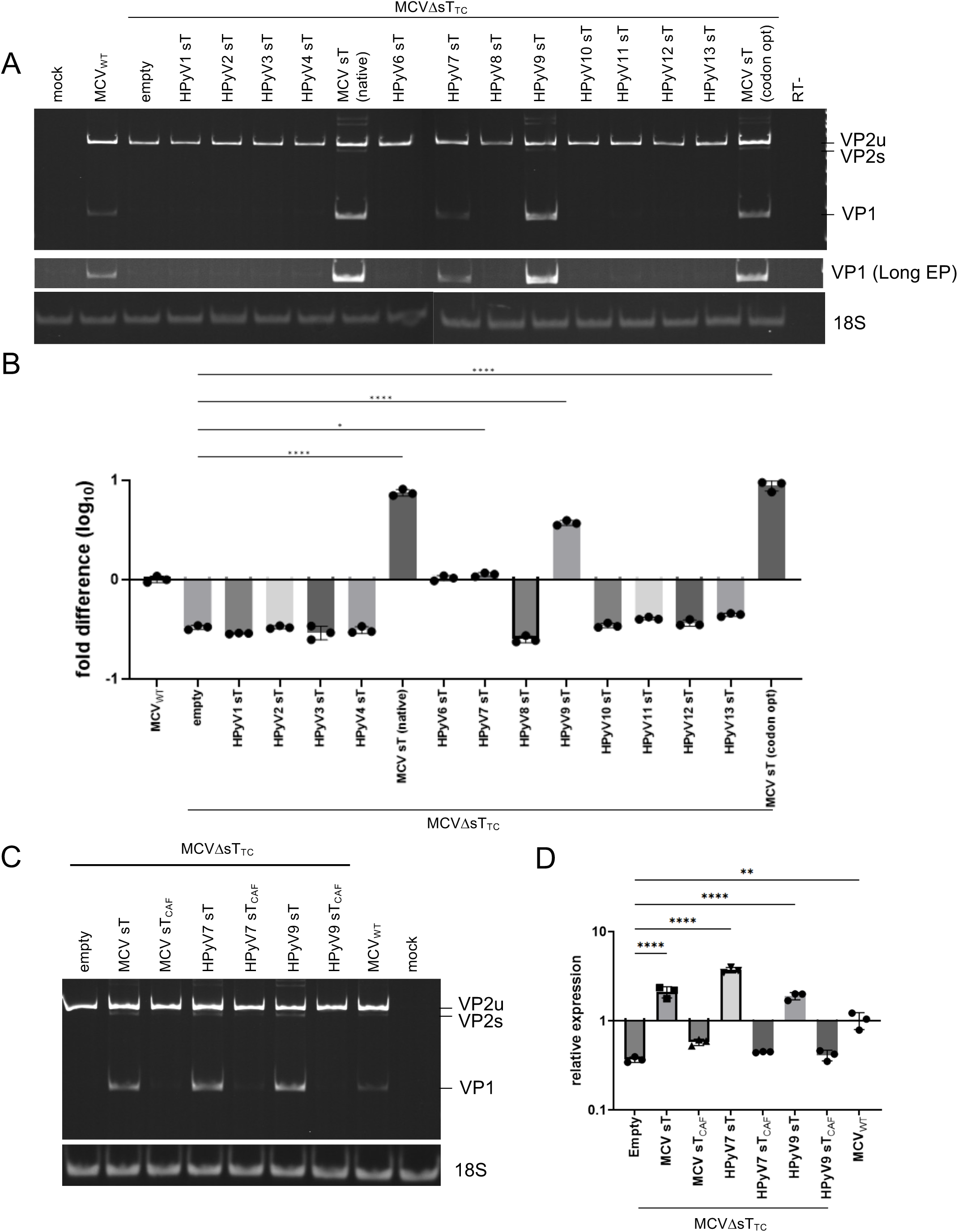
HPyV7 and HPyV9 sT rescue MCV late gene splicing. **(A**) MCV sT function to promote MCV VP1 splicing is conserved in HPyV7 and HPyV9 sT. 293 cells were co-transfected with MCVΔsT_TC_ and various FLAG-sT constructs from human polyomaviruses HPyV1-13. The samples were subjected to RT-PCR. **(B)** Samples in panel **A** were subjected to qRT-PCR analysis. Statistical significance was determined using One-way ANOVA. *, p<0.05; ****, p<0.0001. **(C)** 293 cells were co-transfected with MCV_WT_ alone or MCVΔsT_TC_ together with empty vector, MCV sT, HPyV7 sT, HPyV9 sT, or their respective CAF mutants. The samples were subjected to RT-PCR. **(D)** Samples in **C** were subjected to qRT-PCR. Statistical significance was determined using One-way ANOVA. **, p<0.01; ****, p<0.0001

## Discussion

A less explored question in the polyomavirus life cycle—and more broadly in viruses with compact, replicating genomes—is whether genome replication controls not only transcription but also the maturation of the RNA that it produces. We previously showed that DNA replication is required for sT-induced late-gene transcriptional activation (5). In this study, we found that late gene splicing is also replication-dependent, establishing sT as a direct regulator of replication-coupled RNA processing, distinct from its established role in transcriptional activation. This RNA-processing activity requires the same sT domains that drive viral DNA replication, directly coupling replication status to transcript maturation, and is rescued by sT proteins from HPyV7 and HPyV9. To our knowledge, this is the first demonstration that a polyomavirus small T antigen directly regulates late mRNA maturation, establishing both a replication-coupled mechanism for the early-to-late switch and a new viral model for studying the crosstalk between genome replication and RNA 3′-end processing.

Late genes possess two polyadenylation signals (LpA) (19), leaving a fraction of transcripts that read through both signals to produce longer transcripts. This leaky termination is evident in **Fig. 3C**, where total RNA-seq reads in the late-strand direction from the nuclear fraction show peaks extending past both the LpAs. LpA readthrough may play a role in the MCV life cycle by generating a wraparound pre-mRNA that transcribes through the circular genome, serving as a precursor to mature late mRNA (31,32). Furthermore, miRNAs targeting T antigens are byproducts of LpA readthrough and are suggested to suppress early gene expression and drive the early to-late transition (12,13), whereas in MCV, miRNAs appear to be produced both by readthrough and by an independent, dedicated promoter (25). A similar readthrough-dependent mechanism has been described for MPyV, where both promoters are transcriptionally active from the start of infection, but early transcripts predominate initially. Late transcript levels rise substantially and surpass early transcripts only after DNA replication begins, likely because late genes preferentially use newly replicated DNA as a template (10). In a related but distinct mechanism, Carmichael and colleagues identified leader-to-leader (LL) splicing induced by poly(A) readthrough on a circular DNA virus and demonstrated its critical role in the early-to-late switch of the viral gene expression program (10,33-36).

Our results show that sT drives late-phase RNA maturation—both splicing and 3′-end processing—independent of transcription. RNA-seq has shown that late gene transcription still occurs without sT, albeit at a reduced level, without yielding mature mRNA. DNA replication inhibitors (e.g., aphidicolin) similarly ablate late transcription and maturation, confirming their dependence on replication. However, sT’s polyadenylation-promoting function at LpA may be independent of DNA replication. Unexpectedly, this effect was lost upon LT co-expression, although LT alone had no effect on mScarlet expression (**Fig. 4C**). This loss may reflect substrate saturation, in which the transcriptional output exceeds the cleavage/polyadenylation machinery’s capacity at LpA and increases readthrough despite presumably unchanged sT activity, consistent with reports that readthrough increases once transcriptional demand exceeds processing capacity although further study was required to confirm this hypothesis (37). Indeed, sT-rescue RNA-seq data in MCVΔsT showed that exogenous sT overexpression, together with endogenous LT, overactivates late gene transcription and reduces the relative rate of late mRNA maturation (**Fig. 3C**). Subtle sT protein expression below the immunoblotting detection limit may be sufficient to balance these two activities. sT’s replication-coupled transcriptional activation and independent polyadenylation-promoting functions may normally balance each other. When combined, they overwhelm the machinery, favoring readthrough over termination.

We previously reported that the LTSD mutant of sT localizes to the cytoplasm rather than the nucleus (5). Here, we found that LTSD and the CxCxxC motifs conserved among mammalian polyomaviruses are required for sT nuclear localization, anchoring it to DNA and chromatin. Forcing nuclear entry of these mutants via SV40 NLS attachment failed to restore sT replication, transcription, or RNA processing functions, indicating that chromatin anchoring itself, not nuclear localization per se, is essential. sT’s interactions with chromatin-resident proteins such as EP400 (38) and CBP (5), and its direct CxCxxC-mediated binding to non-methylated CpG islands (39), may explain how sT is physically positioned to couple replicating chromatin to the RNA processing machinery.

The nuclear accumulation of immature wraparound transcripts observed in our study may serve the virus in multiple ways. Because the generation of this pre-mRNA is intrinsically tied to ongoing genome replication, delaying its maturation until sT-dependent processing is complete could function as a checkpoint that couples capsid gene expression to genome copy number, preventing the translation of VP1/VP2 mRNA until sufficient replicated genomes are available for packaging. In addition, prolonged nuclear retention of the unprocessed transcript may extend its availability as a substrate for the microprocessor complex, favoring its processing into T antigen-targeting miRNAs generated by LpA readthrough (25). Supporting a direct role for sT in this process, our previous proximity-labeling (BioID) study identified sT as proximal to DGCR8 (5), a core microprocessor component, raising the possibility that sT facilitates not only polyadenylation of the wraparound transcript but also its processing into viral miRNAs. Together, these observations suggest that sT-mediated maturation of late pre-mRNA may coordinate both the timing of capsid protein expression and the biogenesis of viral miRNAs, which reinforce the early to-late transition.

This replication-coupled RNA processing function appears to be layered onto a broader and more variable role for sT in genome replication. Historically, the importance of sT has been investigated in the animal polyomaviruses SV40 (40) and MPyV (41), whose sT proteins have been shown to indirectly promote viral DNA replication. Among human polyomaviruses, the effect of sT on DNA replication diverges: MCV sT (5,42) and JCV sT (43) promote replication, whereas BKV sT inhibits it (6). The replication-promoting activity of MCV sT depends, in part, on LT stabilization (44), although a stabilization-independent mechanism has also been reported (18). To determine whether this RNA-processing function could be substituted by other HPyV sT, we cloned 13 HPyVs sT for their ability to rescue late gene splicing in MCVΔsT_TC_. Only MCV (HPyV5), HPyV7, and HPyV9 sT — the latter two associated with cutaneous disease: pruritic dyskeratotic dermatosis (45) and severe cutaneous eruptions (46), respectively — reproduced this splicing-rescue phenotype (Fig. 3A), indicating that this activity is not broadly conserved across the polyomavirus family but is instead restricted to a subset of HPyVs. HPyV6 sT, by contrast, restored late gene transcription without rescuing splicing (Fig. 3A-B), reinforcing that transcriptional activation and RNA processing are genetically separable functions of sT. CxCxxC mutagenesis confirmed that this motif is required for splicing rescue by MCV, HPyV7, and HPyV9 sT alike. However, the motif itself is conserved across all 13 HPyV sT proteins tested, indicating that it is necessary but not sufficient to confer this activity. AlphaFold3 comparison of all 13 sT proteins showed an overall conserved fold (**Fig. S7A-B)**, arguing against gross structural divergence as the distinguishing factor and pointing instead to a more local property, such as sequence variation flanking CxCxxC, surface charge, or another feature invisible to global structure prediction, which determines which sT proteins can engage the RNA-processing machinery. Further experiments using whole genome from HPyV6, HPyV7, and HPyV9 instead of MCV are desirable to determine the conservancy of this machinery in other HPyVs.

Host factors that couple sT-driven DNA replication to late RNA processing remain unidentified, representing an important direction for future studies. Because this coupling occurs at the interface between the replisome and RNA processing machinery, strong candidates include replication-associated chromatin factors capable of engaging both the cleavage/polyadenylation and splicing machinery, potentially acting as scaffolds that recruit RNA processing factors to actively replicate viral chromatin. Identifying such factors and determining whether they act locally at the NCCR/late promoter, at the late poly(A) signal, or more broadly across the entire late transcription unit, will be essential for understanding how sT achieves replication-dependent control of RNA maturation.

Genome replication is increasingly recognized as a signal that shapes downstream RNA metabolism and is not merely a prerequisite for it. This study establishes MCV sT as a replication-coupled regulator of late gene RNA processing, directly linking the splicing and polyadenylation of wraparound transcripts to ongoing genome replication, rather than transcription alone. This function, which can be rescued by sT from a subset of related HPyVs, including HPyV7 and HPyV9, extends the known repertoire of polyomavirus sT activities beyond DNA replication and LT stabilization, positioning sT as a central coordinator of the early to-late transition. The apparent balance between sT’s replication-linked transcriptional activation and its independent polyadenylation-promoting activity further suggests that polyomaviruses actively tune, rather than simply tolerate, readthrough transcription inherent to their compact genomes. Defining the host machinery sT engages will be essential to understand how replicating genomes couple duplication to productive RNA maturation.

## Acknowledgement

We thank Dr. Gordon Carmichael for reading our draft manuscript and providing suggestions, and Dr. Patrick Moore and Dr. Yuan Chang for helpful discussion. We also thank Feride Elif Can and Omkar Betsur for their technical assistance, the Health Science Sequencing Core at UPMC Children’s Hospital of Pittsburgh for next-generation sequencing assistance, and Rachel Pekor for administrative support.

## Data availability

R script, Python script, and Bedgraph can be accessed at GitHub (https://github.com/jefferyua-cyber/MCV_splicing.git) and raw RNAseq data is available upon request.

## Supplementary Information

### Supplementary Figure Legend

**Figure S1.**
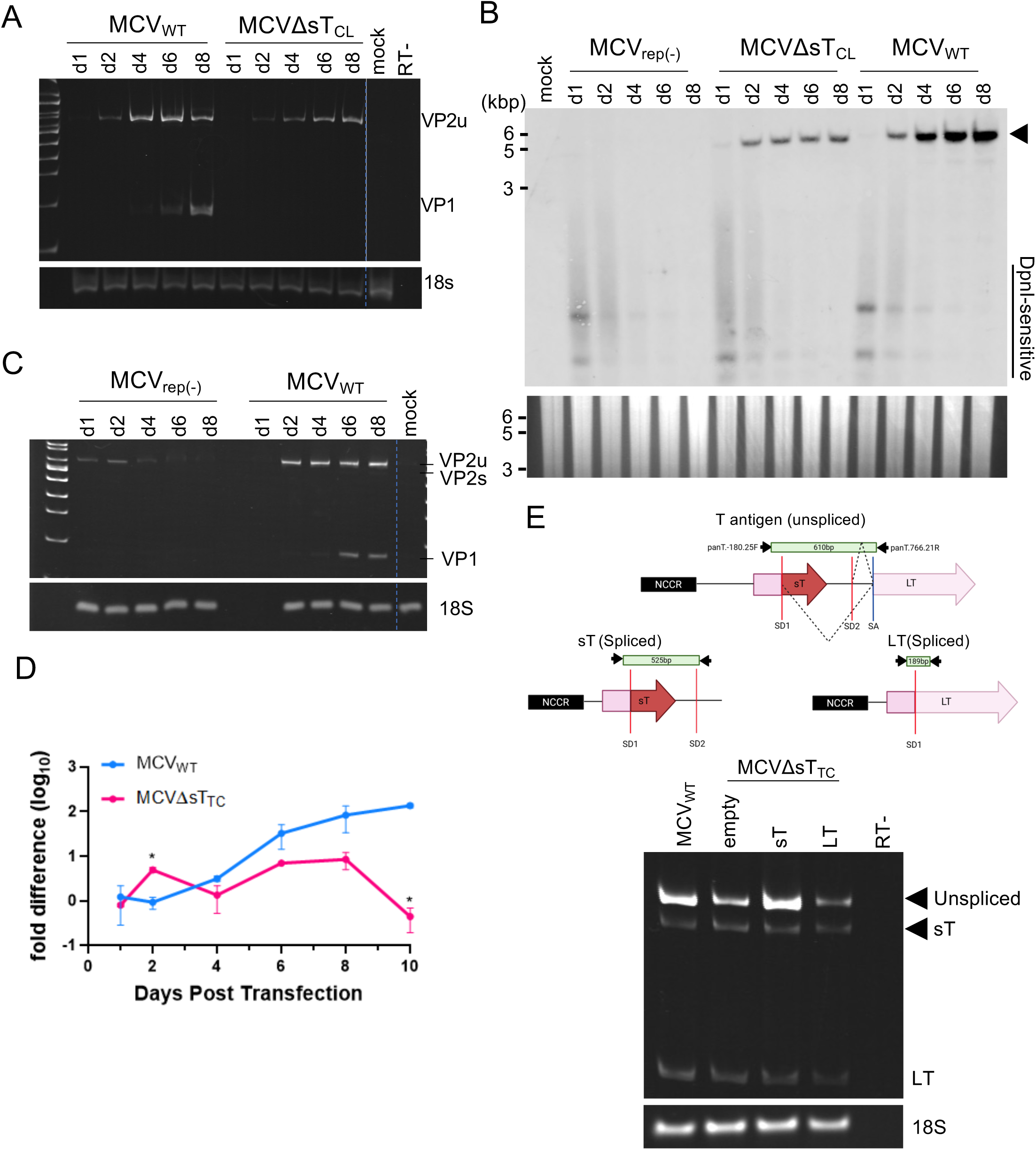
Late gene splicing is affected by sT, but not early gene. **(A)** 293 cells were transfected with MCV_WT_ or MCVΔsT_CL_, cells were collected up to day 8 post-transfection, and total RNA was subjected to random hexamer RT-PCR. Mock and no RT were used as negative controls, and 18S was used as a loading control. **(B)** 293 cells were transfected with MCV_WT_, MCVΔsT_CL_, or MCV_rep(-)_ and collected until day 8. Genomic DNA was extracted, digested with EcoRI and KpnI, and subjected to Southern blotting using an MCV probe. MCV genome was indicated by arrow. **(C)** 293 cells were transfected with MCV_WT_ or MCV_rep(-)_, cells were collected up to day 8 post-transfection, and total RNA was subjected to random hexamer RT-PCR. Mock was used as a negative control, and 18S was used as a loading control. **(D)** Quantitative RT-PCR of samples from Fig. 1B was performed using strand-specific RT-PCR to avoid late gene readthrough contamination, and an early gene-specific primer set was used. **(E)** Schematic representation of the early gene primers used to detect early gene splicing. It produces unspliced T antigen, sT, or LT with 610, 525, and 189 bp size bands. 293 cells were co-transfected with MCV_WT_ or MCVΔsT_TC_ and empty vector, sT, or LT. Total RNA was subjected to strand-specific RT-PCR to detect unspliced T antigen, sT, or LT. No RT was used as a negative control and 18S was used as a loading control.

**Figure S2.**
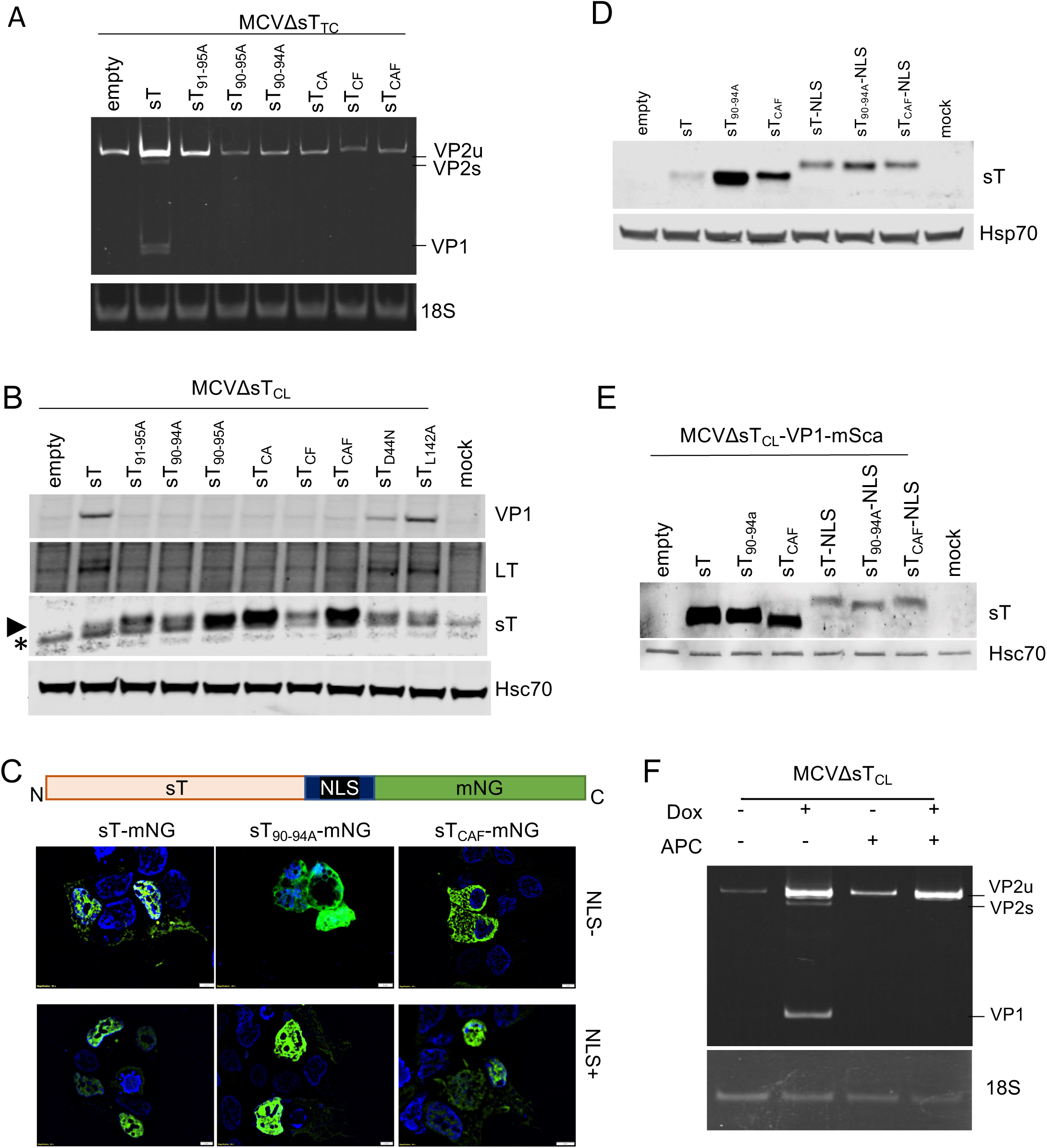
DNA replication and replication domain of sT is required for late gene splicing. **(A)** 293 cells were co-transfected with MCVΔsT_TC_ with empty vector and sT, sT_91-95A_, sT_90-95A_, sT_LTSD_, sT_CA_, sT_CF_, or sT_CAF_, and total RNA was subjected to random hexamer RT-PCR. 18S was used as a loading control. **(B)** 293 cells were co-transfected with MCVΔsT_TC_ or empty vector, sT, sT_91-95A_, sT_90-95A_, sT_LTSD_, sT_CA_, sT_CF_, sT_CAF,_ sT_D44N_, or sT_L142A_ plasmids. Cell pellets were collected, lysed, and subjected to western blotting. The membrane was probed with anti-VP1, LT, sT, or Hsc70 antibodies as loading controls. Target protein bands are indicated by arrows, and non-specific bands are indicated by asterisks, if applicable. **(C-D)** 293 cells were transfected with mNG-tagged empty vector, sT, sT_90-94A_, sT_CAF_, sT-NLS, sT_LTSD_-NLS, or sT_CAF_- NLS. Two days after transfection, the cells were visualized using fluorescence microscopy (panel C) to confirm localization,or were lysed and subjected to western blotting to confirm protein expression (panel **D)**. **(E)** 293 cells were co-transfected with MCVΔST-VP1-mSca and empty vector, sT, sT_90-94A_, sT_CAF_, sT-NLS, sT_90-94A_-NLS, or sT_CAF_-NLS, as shown in Figure 5E-F. The cells were lysed, subjected to western blotting, and probed with an anti-sT antibody to confirm sT expression. Hsc70 was used as the loading control for western blotting analysis. **(F)** 293-TRE-sT cells were transfected with MCVΔsT_CL,_ sT expression was induced using doxycycline, and the cells were treated with APC (5 µg/mL). RNA was extracted and subjected to semi-quantitative RT-PCR.

**Figure S3.**
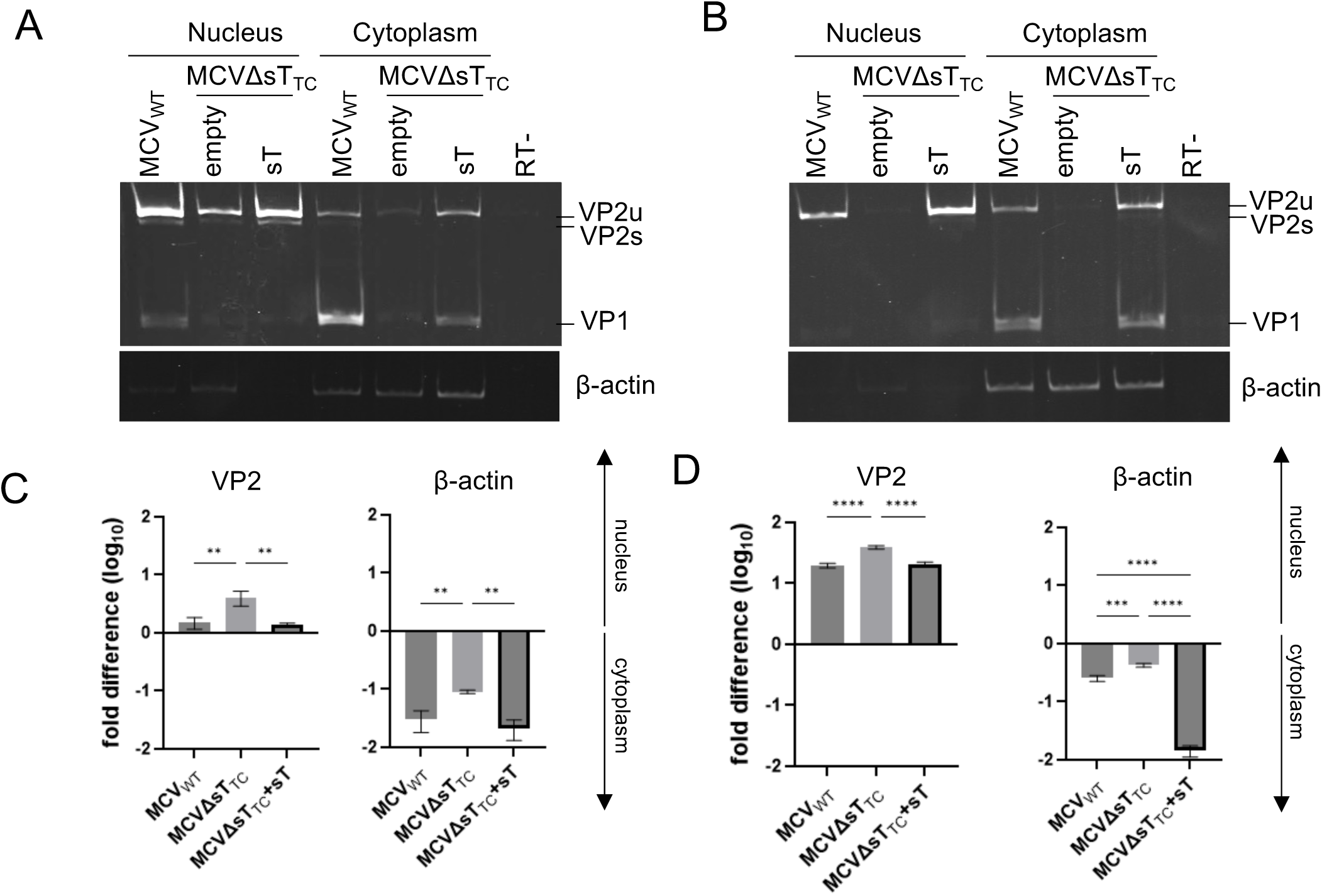
sT is affecting late gene localization. **(A-B)** MCV_WT_ or MCVΔsT_TC_ was transfected into 293-TRE-sT cells (panel **A**) or infected with BJ-hTERT-sT cells (panel **B**). Cells were induced on day one and harvested on day six (panel **A**) or induced on day three and harvested on day seven (panel **B**). Both samples were fractionated to separate the nuclear and cytoplasmic fractions, and total RNA from each fraction was subjected to random hexamer RT-PCR to detect VP1 and VP2 RNA. No RT was used as a negative control, and β-actin was used as a loading control. **(C-D)** qRT-PCR of samples from panels **A** and **B** was performed. The resulting cytoplasmic transcript abundance was normalized against nuclear transcript abundance using the 2^-ΔΔCt^ method to determine localization tendency. Statistical analyses were performed using one-way analysis of variance (ANOVA). **, p<0.01; ***, p<0.001; ****, p<0.0001.

**Figure S4.**
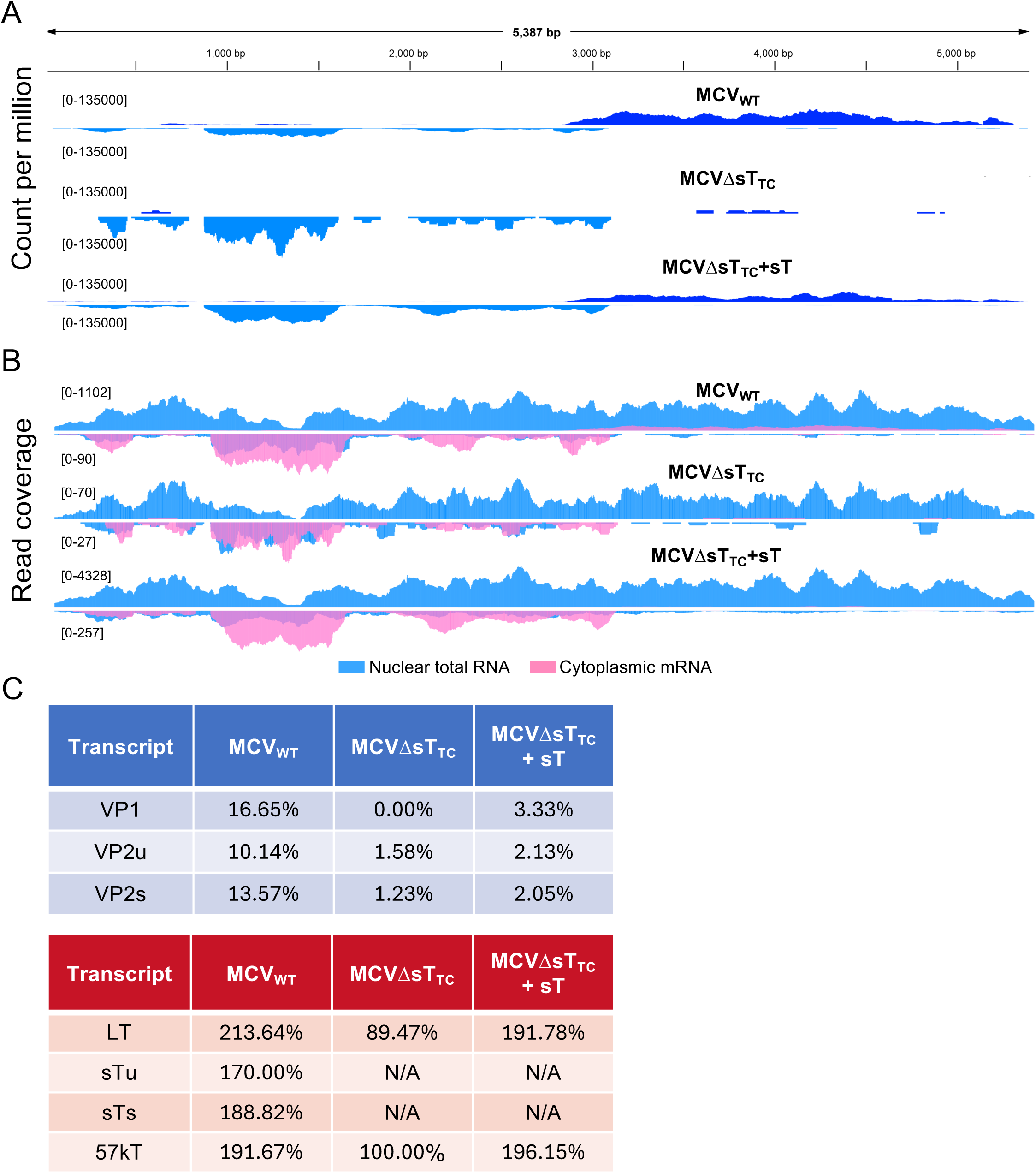
sT deletion hinders early-to-late switch in MCVΔsT_TC_. **(A)** Normalized read coverage of poly(A)-enriched RNA-seq of MCV_WT_ and MCVΔsT_TC_ from the cytoplasmic fraction (related to Figure 2C). 293-TRE-sT cells were transfected with MCV_WT_ or MCVΔsT_TC_ ± sT and induced one day post-transfection. RNA was fractionated between the nuclear and cytoplasmic fractions during extraction. Late gene reads are depicted in the top panel, and early gene reads are depicted on the inverted y-axis in the bottom panel for each sample. Number signifies y-axis range read counts. **(B)** Read coverage of nuclear ribo-depleted total nuclear RNA (blue) and poly(A)-enriched cytoplasmic mRNA (pink) of MCV_WT_- and MCVΔsT_TC_ ± sT-transfected cells. Late genes are depicted in the top panel, and early genes are depicted on the inverted y-axis of the bottom panel. **(C)** Maturation rates of MCV_WT_, MCVΔsT_TC_, and MCVΔsT_TC_ with sT transcripts were determined by dividing the count of poly(A)-enriched RNA-seq with ribo-depleted total RNA and are presented as a percentage.

**Figure S5.**
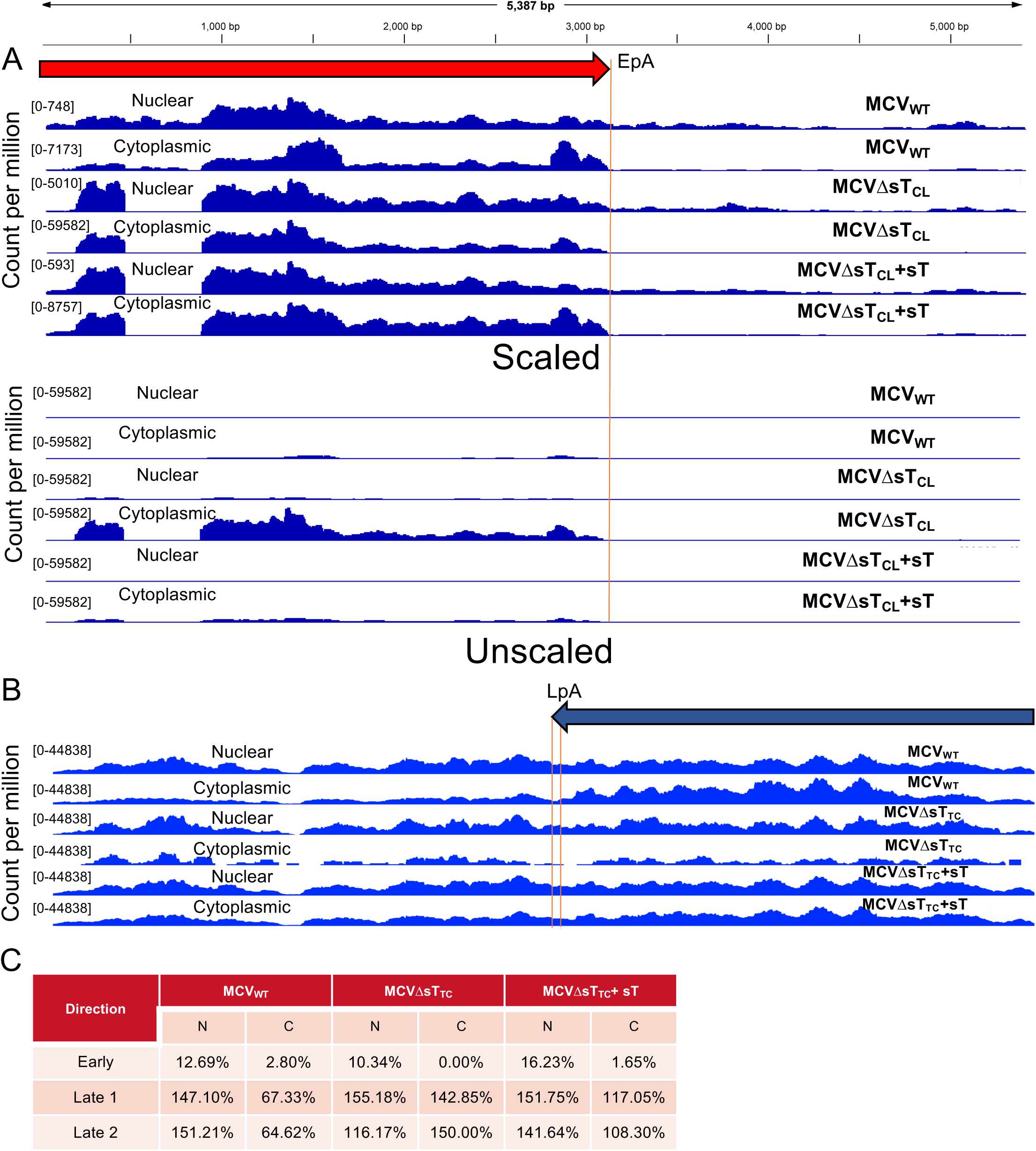
sT is influencing LpA readthrough. **(A)** Normalized read coverage ribo-depleted total RNA-seq of early genes from 293-TRE-ST cells transfected with MCV_WT_ or MCVΔsT_CL_ ± sT from the nuclear and cytoplasmic fractions. **(B)** Normalized read coverage of ribo-depleted total RNA-seq of late genes from 293-TRE-sT cells transfected with MCV_WT_ or MCVΔsT_TC_ ± sT from nuclear and cytoplasmic fractions (related to Fig. 3C). **(C)** The readthrough rate was determined by dividing the transcript count of ribo-depleted total RNA-seq after the respective poly(A) signal (early, late 1, and late 2) by the transcript count before the poly (A) signal and is presented as a percentage. N=nucleus; C=cytoplasm.

**Figure S6.**
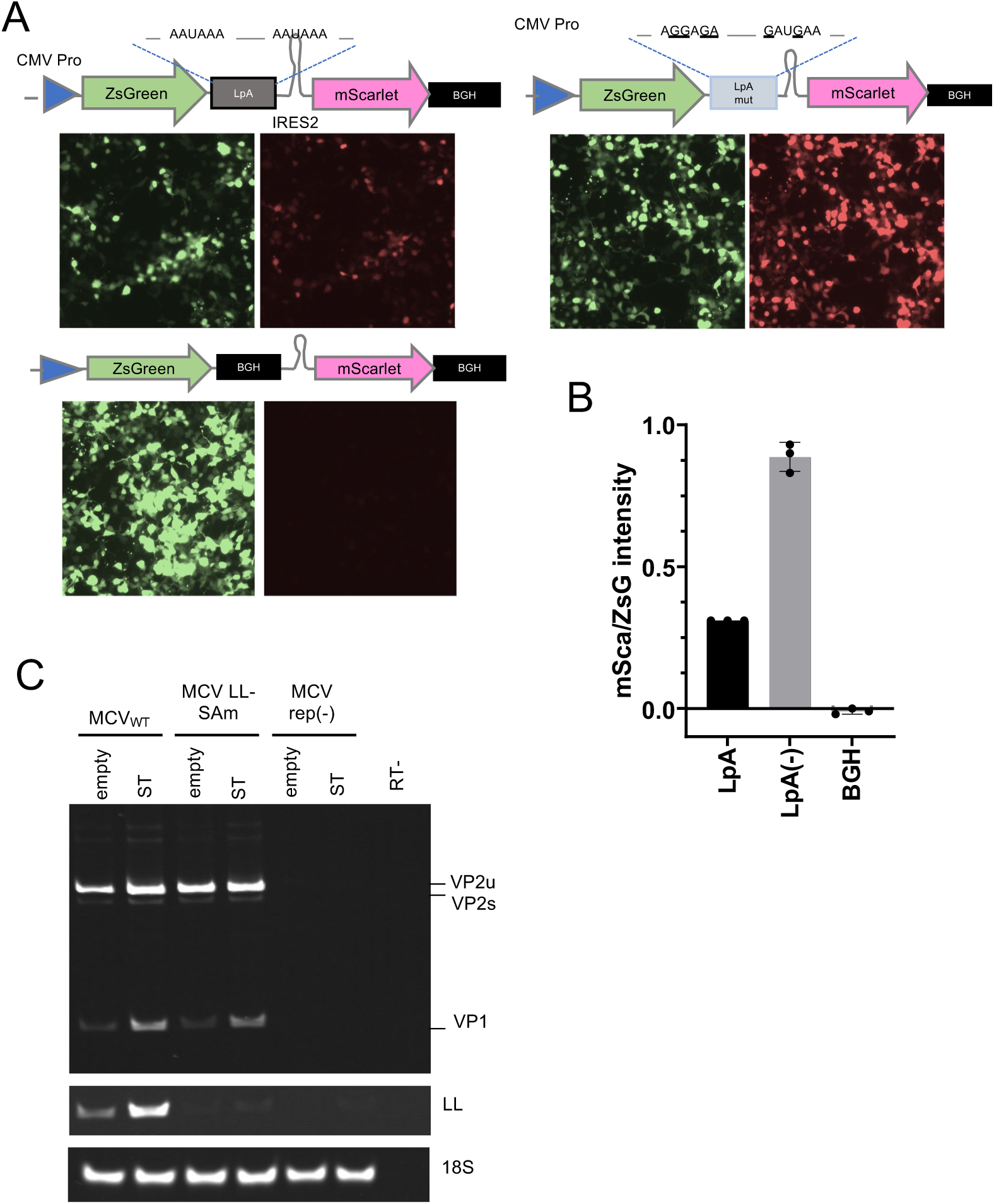
MCV late polyadenylations are leaky. **(A)** Dual reporter system to define LpA readthroughness. The ZsG sequence was followed by various poly(A) (LpA, LpA(-), or BGH), IRES, and mScarlet with BGH poly(A) sequences, as shown in the top panel. 293 cells were transfected with each construct, and on day two post-transfection, ZsG and mScarlet were visualized using a fluorescence microscope. **(B)** Quantification of mScarlet intensity normalized to ZsG. **(C)** MCV_WT,_ MCV LL-SAm, and MCV_rep(-)_ were transfected with empty or sT vectors. Total RNA was subjected to random hexamer RT-PCR. No RT was used as a negative control, and 18S was used as a loading control.

**Figure S7.**
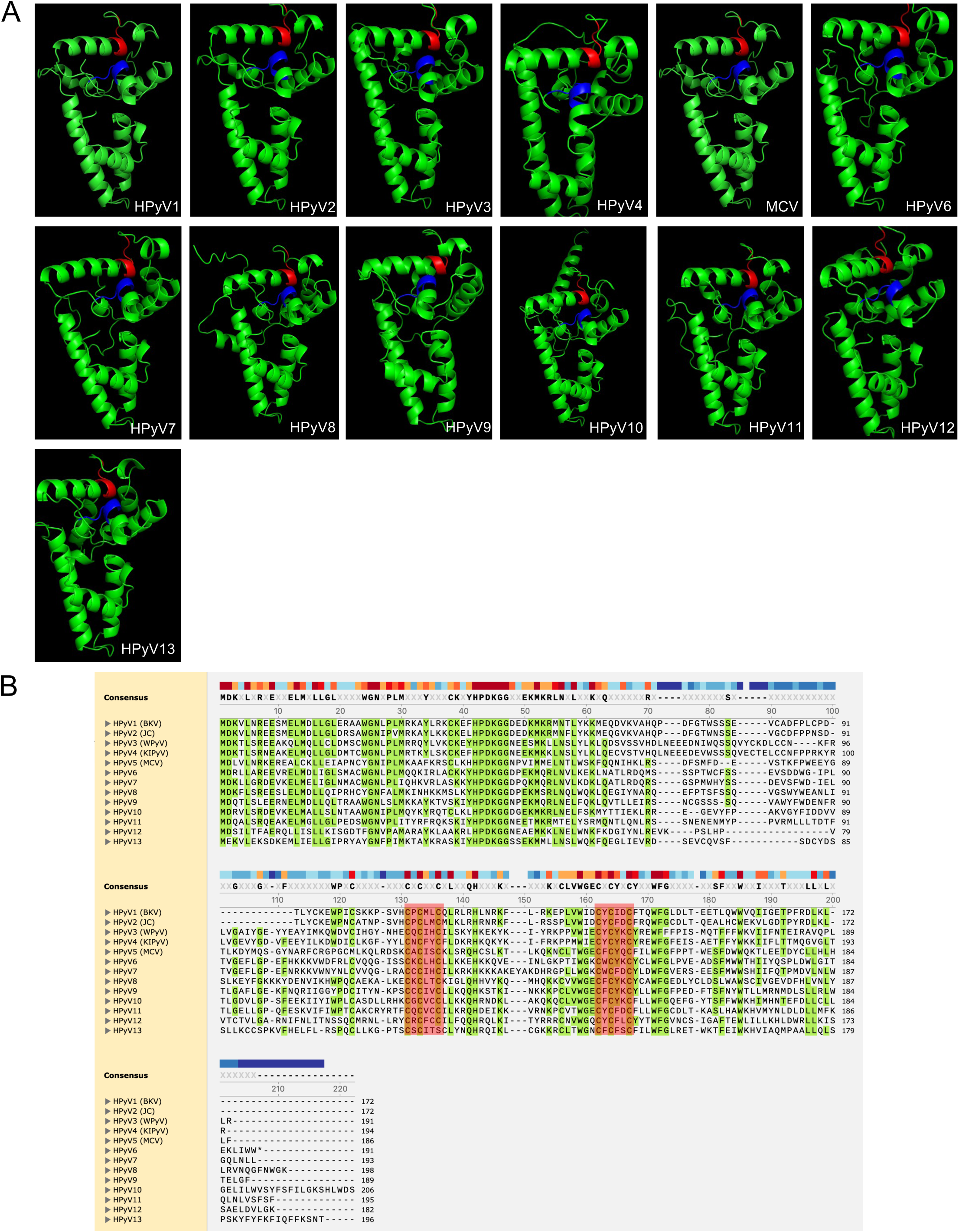
CxCxxC motif is conserved across HpyVs. (A) Human polyomaviruses sT structure prediction using AlphaFold3. Iron-sulfur binding (CxCxxC) motif 1 and 2 were represented in red and blue, respectively. (**B**) Sequence alignment of HPyV sT was performed using ClustalW on SnapGene. The conserved CxCxxC regions are highlighted in red.

